# Broader functions of TIR domains in Arabidopsis immunity

**DOI:** 10.1101/2022.12.23.521769

**Authors:** Pierre Jacob, Junko Hige, Lijiang Song, Adam Bayless, Dor Russ, Vera Bonardi, Farid El-Kasmi, Lisa Wünsch, Yu Yang, Connor R. Fitzpatrick, Brock J. McKinney, Marc T. Nishimura, Murray R. Grant, Jeffery L. Dangl

## Abstract

TIR domains are NAD-degrading enzymes that function during immune signaling in prokaryotes, plants, and animals. In plants, most TIR domains are incorporated into intracellular immune receptors. In Arabidopsis, TIR-derived small molecules bind and activate EDS1 heterodimers, which in turn activate RNLs, a class of cation channel-forming immune receptors. RNL activation drives cytoplasmic Ca^2+^ influx, transcriptional reprogramming, pathogen resistance and host cell death. We screened for mutants that suppress an RNL activation mimic allele and identified a TIR-containing immune receptor, SADR1. Despite functioning downstream of an auto-activated RNL, SADR1 is not required for defense signaling triggered by other tested TIR-containing immune receptors. SADR1 is required for defense signaling initiated by some trans-membrane pattern recognition receptors and contributes to the unbridled spread of cell death in *lesion simulating disease 1*. Together with RNLs, SADR1 regulates defense gene expression at infection site borders, likely in a non-autonomous manner. RNL mutants that cannot sustain this pattern of gene expression are unable to prevent disease spread beyond localized infection sites, suggesting that this pattern corresponds to a pathogen containment mechanism. SADR1 potentiates RNL-driven immune signaling partially through the activation of EDS1, but also partially independently of EDS1. We studied EDS1-independent TIR function using nicotinamide, an NADase inhibitor. We observed decreased defense induction from trans-membrane pattern recognition receptors and decreased calcium influx, pathogen growth restriction and host cell death following intracellular immune receptor activation. We demonstrate that TIR domains can potentiate calcium influx and defense and are thus broadly required for Arabidopsis immunity.

## Introduction

Toll-Interleukin-1 receptor, disease Resistance gene (TIR) domain-containing proteins are conserved from prokaryotes to plants and animals where they regulate immunity and cell death (1). In plants, TIR domains are typically found at the N-termini of Nucleotide-binding Leucine rich repeat immune Receptors (NLRs), a class of intracellular immune receptors triggering a potent immune response called ETI (Effector Triggered Immunity), often associated with host cell death localized to the infection site (2). TIR domains are also encoded as single domain proteins in plants (2). TIR NLRs, hereafter TNLs, are activated upon recognition of pathogen virulence effectors that function to block or dampen immune responses. After effector recognition, TNLs oligomerize to form enzymes that produce a suite of small molecules, including 2’-cADPR, 3’-cADPR, pRib-AMP/ADP, diADPR/ADPR-ATP or 2’,3’-cAMP/cGMP (3–7). pRib-AMP/ADP and diADPR/ADPR-ATP can bind and activate Enhanced Disease Susceptibility 1-Phytoalexin Deficient 4 (EDS1-PAD4) or EDS1-Senescence Associated Gene 101 (EDS1-SAG101) heterodimers, respectively, leading to the recruitment and activation of “helper” NLRs (1, 4, 5, 8). Activated helper NLRs, also termed RNLs due to their N-terminal CC-R domains [RPW8 (Resistance to Powdery Mildew 8)-like Coiled-coil (CC) domain], form Ca^2+^-permeable channels in the plasma membrane, as do some CC-NLRs (hereafter, CNLs) (9, 10). Arabidopsis possesses five active RNLs: Activated Disease Resistance 1 (ADR1), ADR1-like 1 (ADR1-L1), ADR1-L2, N Requirement Gene 1.1 (NRG1.1) and NRG1.2. ADRs and NRGs are partially redundant regulators of immunity and cell death downstream of TNLs (11–13). Ca^2+^ channel blockers and auto-active Ca^2+^ channel mutants indicate that Ca^2+^ influx is necessary and sufficient for Arabidopsis immunity (14).

In a forward genetic screen, we sought to identify new genes required for immunity and cell death activation by RNLs. We found that the TNL *Suppressor of ADR1-L2 1* (*SADR1*) is required for the phenotypes driven by ADR1-L2 auto-activity but is dispensable for other TNL functions. We found that SADR1 regulates defense triggered by the activation of a plasma-membrane pattern recognition receptor and the “runaway cell death” phenotype in the Arabidopsis mutant *lesion simulating disease 1* (*lsd1*), (15). Because these responses involve the perception of extracellular signals, we investigated the requirement for SADR1 and RNLs in the spatial regulation of defense. Virulence effectors delivered to the plant cell from the pathogen *Pseudomonas syringae* pv. *tomato* (*Pst*) DC3000, increase transcriptional defense responses around the infection site. This pattern of host gene expression requires RNLs and SADR1. The loss of defense gene expression on the infection border is associated with the systemic spread of *Pst* DC3000. We found that SADR1 is required for the residual ADR1-L2 auto-activity in the absence of EDS1. These results indicate that SADR1 functions downstream of ADR1-L2 activation partially independently of EDS1 and is thus distinct from the canonical TNL-EDS1-RNL pathway. We tested the requirement for EDS1-independent TIR function in plant immunity using a pharmacological inhibitor of TIR-dependent NADase enzymatic function. We discovered that TIR function is generally required to potentiate immune responses triggered by a plasma-membrane pattern recognition receptor, RNLs and CNLs. Importantly, inhibition of TIR function decreased Ca^2+^ influx resulting from RNLs and CNLs, suggesting that TIR function can generally potentiate Ca^2+^ influx in the context of immune signaling.

## Results

To identify signaling components downstream of ADR1-L2, we screened for mutants able to suppress the auto-immunity associated stunted growth phenotype of an Arabidopsis transgenic line expressing the activation-mimic mutant ADR1-L2 D484V from the native promoter, hereafter ADR1-L2 DV [(16), see Methods for full genotype]. This mutation in the MHD motif is commonly used to mimic NLR activation and can be suppressed in *cis* by P-loop mutations (17–20). ADR1-L2 DV expressing plants exhibit hallmarks of auto-immune signaling: stunted growth, ectopic cell death activation, ectopic salicylic acid accumulation and induction of defense gene expression, including ADR1-L2 itself [Fig. 1 and (16)].

**Figure 1.**
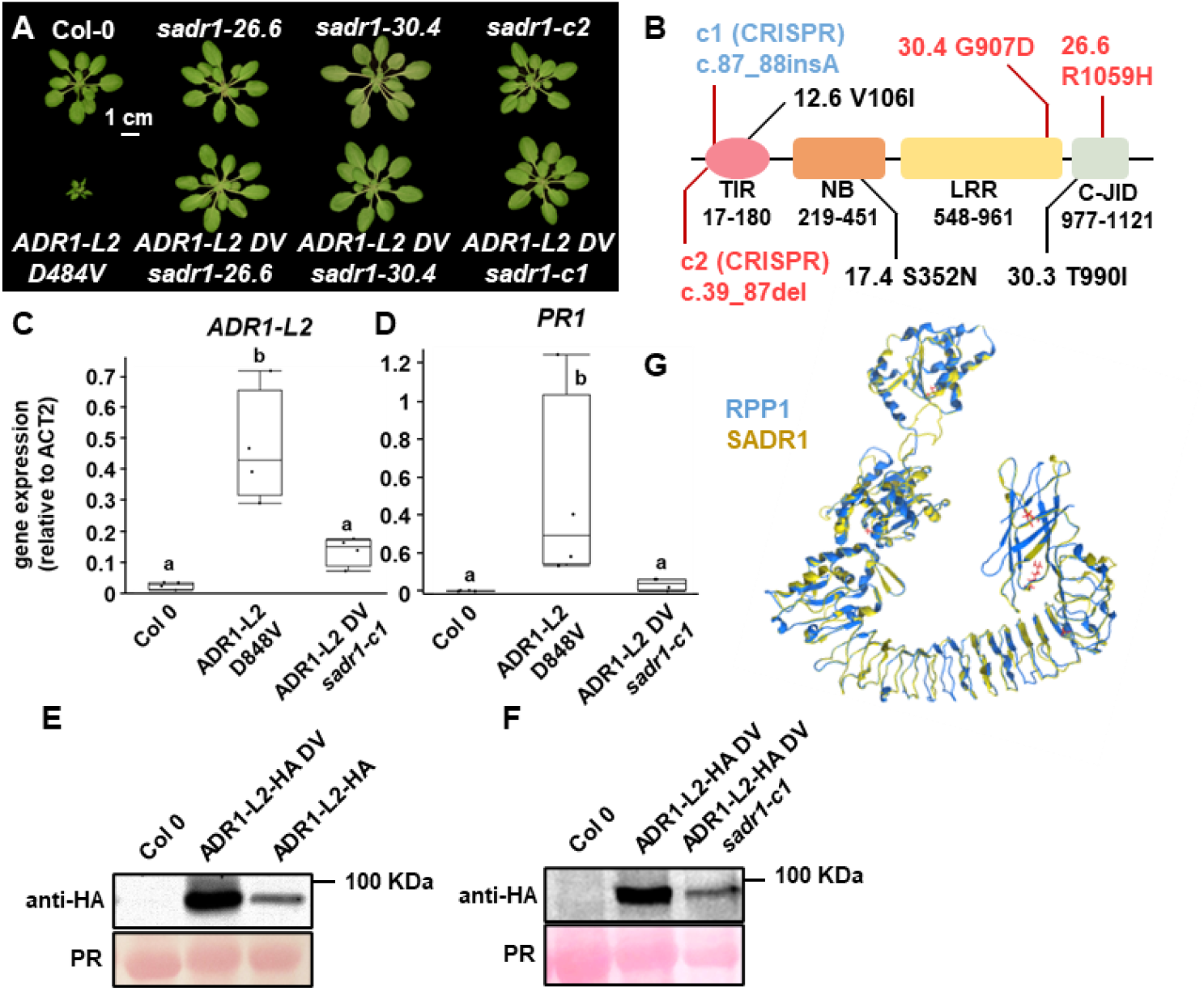
SADR1 is required for the constitutive immunity phenotypes of the activation mimic RNL ADR1-L2 D484V. (**A**) Mutations *sadr1-26.6* and *30.4* fully suppress the stunted growth phenotype of *adr1-l2-4 pADR1:ADR1-L2 D484V* (hereafter ADR1-L2 DV). Introduction of a loss of function mutation in *SADR1* by CRISPR-Cas9 (*sadr1-c1*) suppresses ADR1-L2 DV. (**B**) Schematic representation of SADR1 shows conserved protein domains and the location of mutations identified in the screen or introduced with CRISPR-Cas9. Mutants 12.6, 17.4 and 30.3 are partial suppressors identified in the screen. The fonts indicate the genetic background; blue is *ADR1-L2 DV* and black and red font indicate a wildtype background. (**C**) Suppressed ADR1-L2 DV *sadr1-c1* plants express wild type levels of ADR1-L2 mRNA. Data are from four independent experiment (*N* = 4). Letters indicate statistical significance (T-test, *P* < 0.05). (**D**) *sadr1-c1* suppresses most of the *PR1* expression induced by ADR1-L2 DV. ADR1-L2 DV-induced over-accumulation (**E**) is also suppressed by *sadr1-c1* (**F**). All experiment were performed at least three times. (**G**) SADR1 protein structure modeled onto the TNL RPP1 structure (7CRC). PR: Ponceau red staining.

We identified phenotypically suppressed mutants and performed bulk segregant analysis using suppressed plants from segregating back-crossed F2 populations. Two mutants, 26.6 and 30.4 had mutations in the same gene, located in a genomic region co-segregating with the suppression phenotype in F2 plants (Fig. 1A and B). Whole genome re-sequencing of 32 additional suppressed M3 mutants allowed the identification of three additional mutant alleles of the same gene, 12.6, 17.4 and 30.3 (Fig. 1B). Overall, 5 out of the 39 suppressor mutants identified were affected in this gene (the others will be described elsewhere), which we consequently named *Suppressor of ADR1-L2 1* (*SADR1*, AT4G36150). RNA-sequencing showed that mutations 26-6 and 30-4 suppressed the vast majority of ADR1-L2 DV-driven gene expression changes (*SI Appendix*, Fig. S1, Table S1). We created a *sadr1-c1* loss of function allele in ADR1-L2 DV expressing plants using CRISPR-Cas9 (c.87_88insA, leading to a frameshift after Q29; Methods). The *sadr1-c1* mutation suppressed the ADR1-L2 DV stunted growth phenotype and constitutive expression of *Pathogenesis Related 1* (*PR1*). Importantly, *ADR1-L2 DV* mRNA and protein levels were reverted to wildtype levels (Fig. 1C, D, E and F). This demonstrates that SADR1 is required for ADR1-L2 DV self-amplification, an important feature of RNL signaling (16, 21).

Surprisingly, *SADR1* encodes a TNL, homologous to Recognition of *Peronospora parasitica* 1 (*RPP1*, Fig. 1G; *SI Appendix*, Fig. S2). *SADR1* is physically located next to another TNL (*SADR1-Paired 1*; AT4G36140), in a head-to-head configuration similar to the sensor/executor TNL pair *RRS1/RPS4* (Resistance to *Ralstonia solanacearum* 1 / Resistance to *P. syringae 4; SI Appendix*, Fig. S2). However, CRISPR-derived *SADR1-P1* loss of function allele *sadr1-p1-c1*, (c.247_332del), did not modify ADR1-L2 DV auto-activity (*SI Appendix*, Fig. S3). Overall, these results indicate that the genetically paired TNL SADR1 is required for the ADR1-L2 DV activation mimic phenotype.

We characterized the function of SADR1 in defense. We generated a *sadr1-c2* loss of function mutant (c.39_88del leading to a frameshift after V12) with CRISPR-Cas9 in the wild-type Col-0 background (Fig. 1B). We observed that SADR1 was not required for basal resistance to the virulent pathogen *Pst* DC3000, or to the avirulent strains *Pst* DC3000 *AvrRpt2* or *Pst* DC3000 *AvrRps4*, which activate the CNL RPS2 and the TNL-pair RPS4/RRS1 immune receptors, respectively (Methods). As a control, we showed that the RNL-defective *helperless* quintuple mutant (11, 13) was indeed more susceptible to infection than Col-0 in each of these situations (*SI Appendix*, Fig. S4, A to C).

We investigated the possibility that loss of SADR1 could be obscured by redundant RNL signaling. We compared *adr1 adr1-l1* and *adr1 adr1-l1 sadr1-c2* to *adr1 adr1-l1 adr1-l2* mutants during TNL-driven immunity following challenge with *Pst* DC3000 *AvrRps4* (*SI Appendix*, Fig. S4D).

ADR1s were required for full bacterial growth restriction in these conditions, as seen with the 10-fold increase in pathogen growth in *adr1 adr1-l1 adr1-l2* compared to Col-0. ADR1-L2 RNL function to ‘help’ RPS2, which was required for approximately half of this growth restriction, was not affected by *sadr1-c2* loss of function (*SI Appendix*, Fig. S4D). Resistance to *Hyaloperonospora arabidopsidis* isolate Cala2, which activates the TNL RPP2, was also not affected by *sadr1-c2* (*SI Appendix*, Fig. S4E and F). In addition, SADR1 was not required for the auto-active phenotype of the *snc1* TNL allele [*SI Appendix*, Fig. S4G and H, (22)]. These results indicate that SADR1 is not required for RNL-driven defense against virulent or avirulent bacteria, at least for the TNL functions we measured.

RNLs are also required for some responses to Pathogen-Associated Molecular Patterns (PAMPs) (23–25). We tested PAMP response in *sadr1-c2* (Col-0 background) using NLP20 [necrosis and ethylene-inducing peptide 1 (Nep1)-like proteins (NLPs)], a widespread PAMP (26). Pre-treatment with NLP20 24h before challenging the plants with *Pst* DC3000 primed defense responses to subsequent inoculation with *Pst* DC3000 in both Col-0 and *adr1-l2* but not in *sadr1-c2* or *helperless* plants (Fig. 2A; Methods). Therefore, SADR1 is required for NLP20-driven defense priming. We did not observe a SADR1 requirement for flg22-priming.

**Figure 2.**
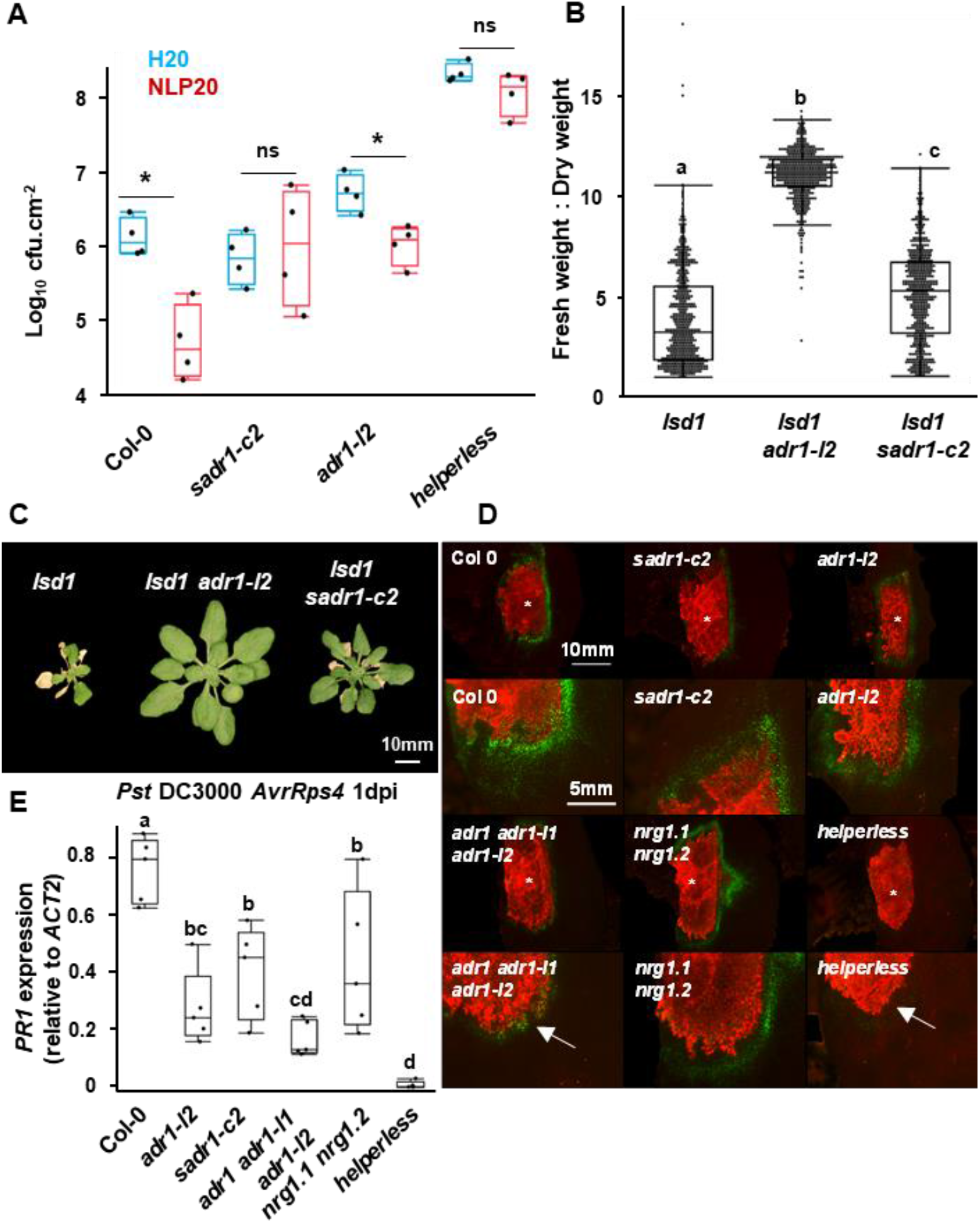
SADR1 is required for NLP20/RLP23 signaling, contributes to *lsd1* runaway cell death, and regulates *PR1* expression around the infection site. (**A**) SADR1 is required downstream of NLP20/RLP23. Plants were challenged with *Pst* DC3000 EV 24 hours after water or NLP20 1μM treatment (*N* = 4) (45). (**B**) SADR1 is partially required for *lsd1* runaway cell death. Fresh:dry weight ratio measurements indicating the proportion of dead tissues two weeks after induction of runaway cell death with 300μM BTH. Data are from six independent experiments (*N* > 70). **(C)** Representative pictures of plants in (B). (**D**) SADR1 and RNLs are required for *PR1* expression at the margin of infection sites. Representative pictures of pPR1:YFP^NLS^-expressing leaves of the indicated genotype infected with *Pst* DC3000 *AvrRps4* mCherry (OD = 0.2) at 24hpi. Notably, *adr1 adr1-l1 adr1-l2* and *helperless* mutants cannot induce strong *PR1* expression on the infection border (white arrows). See *SI Appendix*, Figure S5. (**E**) *PR1* expression on the margin of the infection site 24h after infection with *Pst* DC3000 *AvrRps4* (*N* = 4). Data presented in (E) are from 5 independent experiments. Letters indicate statistical significance [(A and B) ANOVA with post hoc Tukey, (E) T-test, *P* < 0.05].

ADR1-L1 and ADR1-L2 are also required for “runaway cell death”, the superoxide-driven self-perpetuating cell death observed in *lesion simulating disease 1* (*lsd1*) (15, 25). To test if SADR1 mediates runaway cell death, we treated four-week-old plants with BTH (Benzothiadiazole), a salicylic acid analog (27) which triggers runaway cell death in *lsd1*. After two weeks, we measured fresh and dry weight of the BTH-treated plants to estimate the extent of cell death induction (Methods). As expected, *lsd1* displayed extensive lesions covering most or all of the plant, which resulted in a very low fresh:dry weight ratio in *lsd1*, compared to the suppressed *lsd1 adr1-l2* phenotype [Fig. 2B and C;(25)]. *lsd1 sadr1-c2* exhibited an intermediate phenotype, indicating that SADR1 contributes positively to runaway cell death (Fig. 2B and C). Overall, SADR1 is not required for TNL signaling, but is involved in PAMP signaling and *lsd1* runaway cell death.

The *lsd1* runaway cell death phenotype is non-cell autonomous because the induction of self-perpetuating cell death depends on the proximity and perception of a dead or dying cell (15). PRR signaling also involves non-autonomous relay of defense gene activation in neighboring cells (28). Similarly, damage-associated molecular patterns (DAMPS) trigger calcium-dependent defense responses in surrounding tissue (29). Consistent with this, expression of *PR1* occurs in the area surrounding the cells undergoing cell death during NLR-mediated immune responses (30, 31). Using reporter plants (31) expressing YFP^NLS^ under the control of the *PR1* promoter, we reproduced and extended these observations. We used mCherry-tagged bacteria and observed that inoculation with either *Pst* DC3000 EV (virulent), *Pst* DC3000 *AvrRpt2* or *Pst* DC3000 *AvrRps4* (activating the CNL RPS2 or the TNL RPS4, respectively) induced a pattern of *PR1* expression at the border of the infection site (*SI Appendix*, Fig. S5). This pattern did not result from inhibition of *PR1* expression in the infection zone by coronatine (*SI Appendix*, Fig. S5), a pathogen-derived phytotoxin and Jasmonic Aacid mimic known to antagonize SA signaling and inhibit *PR1* expression (32). The pattern of *PR1* expression was not observed in plants challenged with *Pst* DC3000 *hrcC*, which cannot deliver virulence effectors, consistent with effector-dependent defense inhibition (*SI Appendix*, Fig. S5). At 6 hpi, *PR1* promoter activity appeared to be enhanced in NLR-activating inoculations compared to *Pst* DC3000 *hrcC*, suggesting that NLR signaling increased defense around the infection site (*SI Appendix*, Fig. S5A). At 24 hpi, bacterial growth led to a visible mCherry signal, largely non-overlapping with YFP positive areas defining *PR1* expression (*SI Appendix*, Fig. S5B). Overall, cells expressing *PR1* are likely not subjected to effector-driven defense inhibition and are spatially separated from the bacteria. These results suggest that NLR signaling relays defense gene activation in areas devoid of type III effectors, possibly through DAMP activation or reactive oxygen-based signaling (15, 29)

We next investigated whether RNLs and SADR1 were involved in spatial regulation of defense. We infected Col-0 *pPR1:YFP^NLS^* reporter plants mutated in *SADR1* or *RNLs* with a high concentration inoculum (OD_600_ = 0.2) of *Pst* DC3000 *AvrRps4* mCherry to activate the TNL RPS4. Fluorescence observation and qPCR quantification of *PR1* mRNA indicated that both SADR1 and RNLs (ADR1s in particular) regulate defense at the borders of infection sites (Fig. 2D and E). These results suggest that SADR1 and RNLs mediate defense gene expression at the borders of infection sites.

Activating defense around the infection area could serve to prevent the systemic propagation of pathogens. To test this hypothesis, we inoculated half-leaves with *Pst* DC3000 EV and isolated non-infiltrated tissues from the same leaves after ten days (Fig. 3; Methods). We found that only a very low level of *Pst* DC3000 propagates to the non-infiltrated side of the leaf in Col-0 (Fig. 3B). Similar levels were found in *sadr1-c2* and *nrg1.1 nrg1.2* mutants. However, we observed a dramatic increase in the spread of disease symptoms and bacterial growth in non-infiltrated tissues in *adr1 adr1-l1 adr1-l2* and *helperless* plants at 10 days post inoculation. *ADR1* defective plants exhibited systemic disease symptoms after four weeks, including reduced growth, anthocyanin accumulation and systemic lesions (Fig. 3D). Overall, RNLs induce defense gene expression at the infection site borders and *ADR1s* in particular are required to limit systemic *Pst* DC3000 propagation and disease spread from a localized infection event. *SADR1* and *NRG1s* contribute to this disease resistance mechanism. These results suggest that the selective activation of ADRs (4, 5) is most relevant for this phenotype.

**Figure 3.**
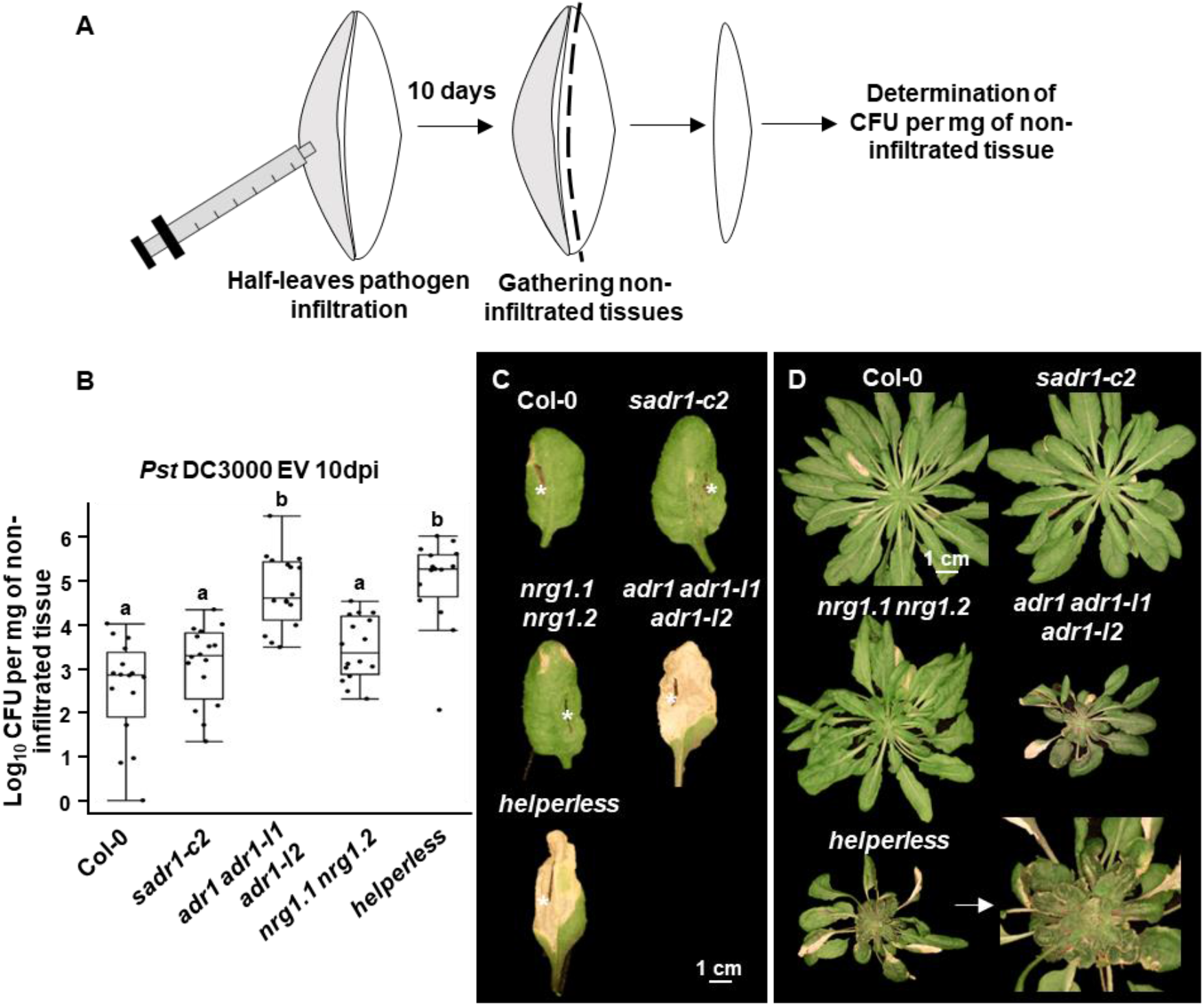
ADR1s limit *Pst* DC3000 propagation and prevent systemic disease from localized infections. (A) Schematic representation of the experimental procedure used in (B) to measure the extent of pathogen propagation *in planta*. (**B**) Bacterial growth at 10dpi in non-infiltrated tissues (*N* = 4, ANOVA with post hoc Tukey, *P* < 0.01). (**C**) Representative pictures of leaves infiltrated with *Pst* DC3000 EV on one half (marked with the white asterisks), at 10dpi. Leaves of the *adr1 adr1-l1 adr1-l2* triple mutant and RNL-free *helperless* plants exhibit expanding lesions into un-infiltrated tissues. (**D**) Representative pictures of plants infiltrated with *Pst* DC3000 on four half leaves, at 28dpi. The *adr1 adr1-l1 adr1-l2* triple mutant and *helperless* plants show systemic disease symptoms.

SADR1 is required for ADR1-L2 DV activation mimic phenotypes, but it is not required for either RPS4, RPP2 or the *snc1* auto-activity TNL phenotypes (Fig. S4). TNLs regulate immunity by activating ADR and NRG RNLs through the selective TIR ligand-bound forms of the EDS1-PAD4 or EDS1-SAG101 heteromeric complexes, respectively (4, 5). We sought to understand why SADR1 TNL activity would be required in a context where an RNL, ADR1-L2, is already active. We hypothesized that SADR1 could be amplifying the defense signal initiated by ADR1-L2 DV in a positive feedback loop, as evidenced by the expression data in Figure 1. Different *eds1* loss of function alleles differentially affect the ADR1-L2 DV auto-immune phenotype (16, 33). We repeated these observations, but with the “clean” CRISPR deletion *eds1-12* allele (34). We found that *eds1-12* only partially suppresses the ADR1-L2 DV stunted growth phenotype (Fig. 4A and B). In contrast, the *sadr1-c1* allele fully suppressed ADR1-L2 DV-driven stunted growth and defense priming (Fig. 4). These results define an EDS1-independent potentiation of ADR1-L2 DV activity by SADR1 that is retained in *ADR1-L2 DV eds1-12* plants (where ADR1-L2 DV is expressed from the native promoter; see *SI Appendix* Methods). We next questioned whether this potentiation of ADR1-L2 DV activity was specific to SADR1. Introgression of the auto-active *snc1* TNL into the fully suppressed *ADR1-L2 DV sadr1-c1* background also restored some ADR1-L2 DV activity, suggesting that potentiation of ADR1-L2 DV can also be provided by at least this additional auto-active TNL (*SI Appendix*, Fig. S6). Therefore, SADR1 can potentiate ADR1-L2 DV activity independently of EDS1 and this function may be shared by other active TIR-containing proteins.

**Figure 4.**
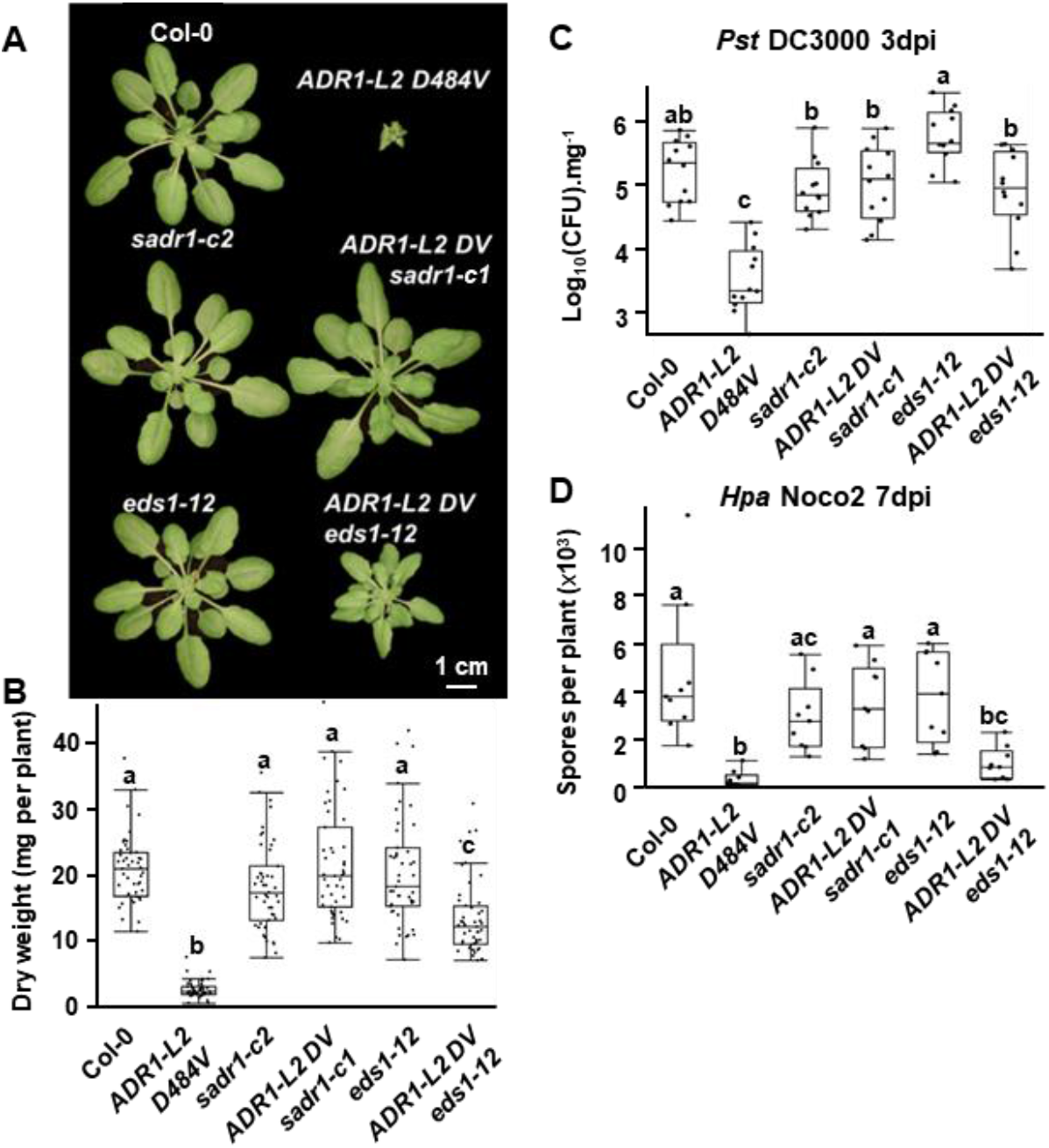
SADR1 potentiates residual ADR1-L2 D484V activity independently of EDS1. (**A**) Representative pictures of 6-weeks old plants with the genotypes indicated above. ADR1-L2 D484V-driven growth inhibition (*N* = 48) (**B**), defense against *Pst* DC3000 (*N* = 12) (**C**), and resistance to *Hpa* isolate Noco2 (*N* = 9) (**D**) are fully suppressed by *sadr1-c1* and partially by *eds1-12*. Data are from three independent experiments. Letters indicate statistical significance (ANOVA with post hoc Tukey, *P* < 0.05).

Arabidopsis possess a large repertoire of TIR-containing proteins (2), many of which are transcriptionally induced during the initial steps of defense signaling. TNL over-expression is often sufficient to trigger immune responses (24). Consequently, evaluating the contribution of potential EDS1-independent-TIR signaling to defense would benefit from an inhibitor of TIR enzymatic function. The first step of plant TIR enzymatic pathway involves cleavage of NAD^+^ into nicotinamide (NAM) and ADPR (35). Interestingly, high concentrations of NAM (50mM) inhibit the NADase activity of the mammalian CD38 NADase and have been deployed in Arabidopsis tissues to inhibit cADPR accumulation, suggesting that NAM could inhibit plant TIR NADase activity (36, 37).

We looked for a readily measurable bio-indicator of TIR enzymatic activity *in planta* because it is difficult to detect the TIR-derived molecules that are the signaling ligands for EDS1-dependent heteromers (4, 5). 2’/3’-RA, [2’-O-β-D-ribofuranosyladenosine or 3’-O-β-D-ribofuranosyladenosine] nucleotide metabolites similar to 2’/3’cADPR but lacking the pyrophosphate groups, accumulate during TIR-dependent plant immune responses *in planta* (7, 38, 39). We transiently expressed in *N. benthamiana* leaves either active full length TNLs or TIR domains fused with SARM1 oligomerization domain, [the SAM domain, which enhances TIR activation; (35)]. We used the corresponding TNLs or TIR domains rendered inactive by mutation of their respective catalytic glutamic acid residues as negative controls (*SI Appendix*, Fig. S7A; Methods). We found that active TNLs or TIR-SAM domain fusions reliably induced the accumulation of 2’/3’-RA, [*SI Appendix*, Fig. S7B; (38)]. This accumulation was dependent on the conserved catalytic glutamic acid in all cases. Interestingly, a SADR1 TIR-SAM domain fusion induced a very small and inconsistent accumulation of 2’/3’-RA suggesting it may act differently than RPP1, RPS4 or BdTIR (*SI Appendix*, Fig. S7B). However, overall, 2’/3’-RA is a reliable and readily measured bio-indicator of TIR enzymatic activity.

We then tested the impact of 50mM NAM treatment on 2’/3’-RA accumulation during pathogen infection. We infected plants with *Pst DC3000ΔhopAM1-1, hopAM1-2*, lacking both copies of the active TIR mimic type III effector *hopAM1* [to avoid HopAM1-produced NADase products; Methods; (39, 40) and also expressing, or not, *AvrRps4* to induce TNL RPS4 activity. We then detected 2’/3’-RA with LC-MS/MS at 12h post infiltration (*SI* Appendix, Methods). An increase in 2’/3’-RA was detected in plants infected with *Pst DC30000ΔhopAM1-1, hopAM1-2 AvrRps4* but not in plants treated with *Pst DC3000ΔhopAM1-1, hopAM1-2*. This result indicates that TNL RPS4 activation leads to 2’/3’-RA accumulation *in planta* (*SI Appendix*, Fig. S8). Consistent with previous reports, the 2’/3’-RA increase was enhanced in *eds1* plants, likely due to the absence of cell death induction (41). Co-treatment with 50mM NAM inhibited TNL RPS4-dependent 2’/3’-RA accumulation *in planta* (*SI Appendix*, Fig. S8). These results are consistent with the hypothesis that 50 mM NAM inhibits TIR enzymatic activity *in planta*.

We therefore used NAM treatment to evaluate the contribution of TIR enzymatic function to defense. TIR activity and subsequent EDS1-dependent immune signaling contributes not only to ETI activated by TNL receptors, but also to basal defense and consequent growth restriction of *Pst* DC3000 EV. This is because basal defense responses include “weak” ETI, at least some of which is likely to be TNL- and EDS1-dependent (42, 43). Also, as noted above, defense responses include transcriptional up-regulation of many TIR domain-encoding genes which could boost immunity via production of TIR enzymatic products to functional levels (1, 24, 44, 45).

We treated Col-0, *sadr1-c2* and *helperless* plants with NLP20, with or without co-inoculation of 50mM NAM to test general TIR function. NAM inhibited *PR1* induction following NLP20 treatment in Col-0 and suppressed it in *sadr1-c2*, which we demonstrated above is required for full NLP20-dependent signaling (Fig. 5A). We then tested the impact of 50mM NAM on defense against virulent and avirulent *Pst* DC3000. NAM significantly inhibited resistance against *Pst* DC3000 EV and *Pst* DC3000 *AvrRps4* in Col-0, but not in *eds1*, consistent with NAM inhibiting an EDS1-dependent defense pathway (Fig. 5B and C). We noted that NAM co-treatment inhibited pathogen growth in otherwise hyper-susceptible *eds1* plants. We therefore tested the impact of NAM on bacterial growth in minimal (MS) or rich culture medium (LB) and found that NAM was also bacteriostatic, potentially explaining why NAM had a negative impact on bacterial growth in *eds1* plants (*SI Appendix*, Fig. S9). These results collectively indicate that the impact of NAM on plant defense as measured by bacterial growth is likely to be underestimated. Overall, NAM inhibits EDS1-dependent defenses.

**Figure 5.**
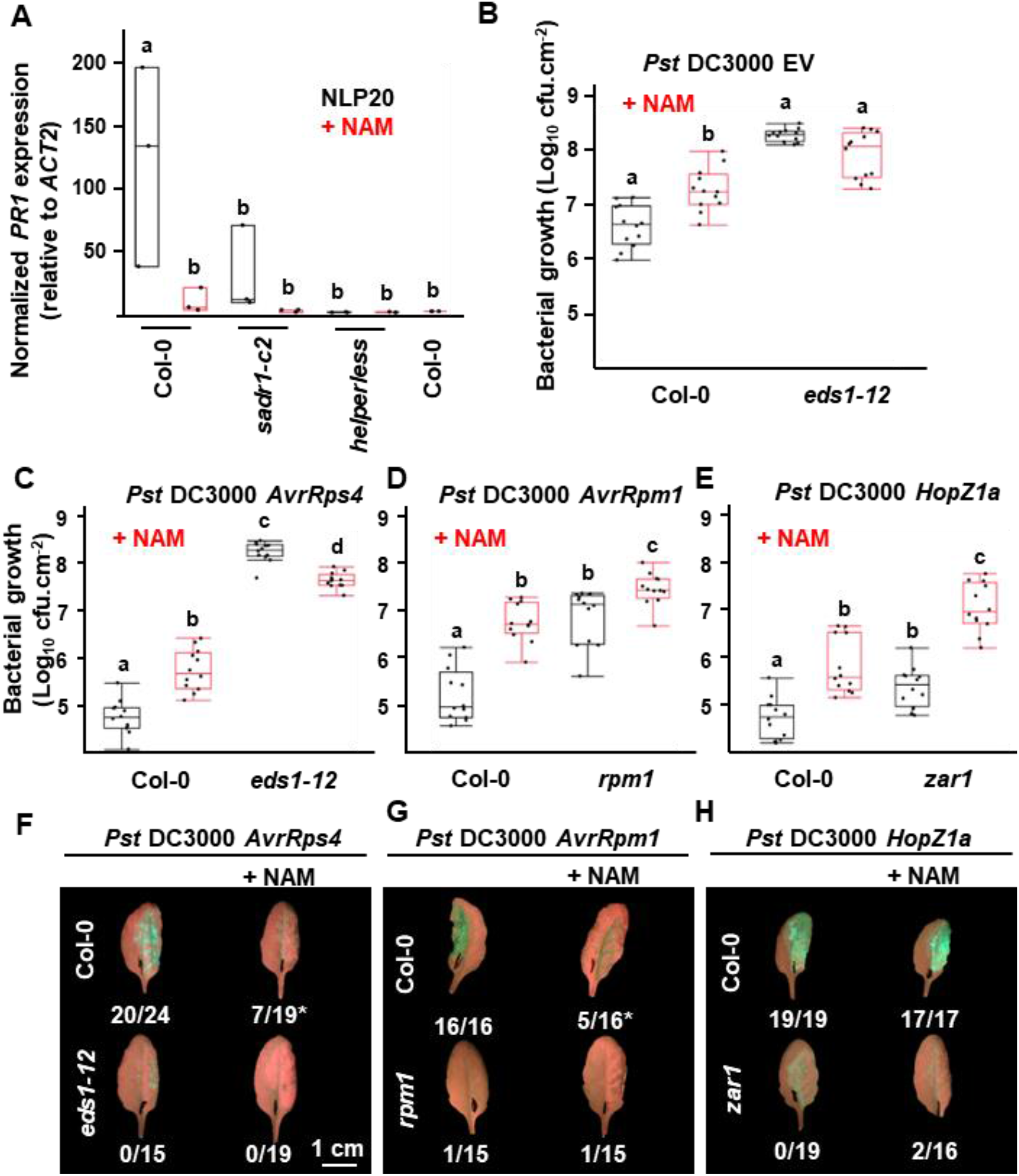
Inhibition of TIR enzymatic activity with NAM regulates NLP20 response, as well as defense and cell death resulting from NLR activation. (**A)** 50mM NAM treatment inhibits *PR1* expression following NLP20 treatment and defense against virulent *Pst* DC3000 EV (*N* = 3) (**B**), avirulent *Pst* DC3000 *AvrRps4* (**C**), *AvrRpm1* (**D**) and *HopZ1a* (**E**), (*N* = 12). NAM treatment also delays cell death induction after inoculation of *Pst* DC3000 (OD_600_ = 0.2) expressing *AvrRps4* (**F**) or *AvrRpm1* (**G**) but not *HopZ1a* (**H**). Numbers indicate the number of HR+ leaves. * Loss of turgor was observed in some leaves that did not exhibit auto-fluorescence characteristic of HR cell death. Data from (A) to (E) are from three independent experiments and from one representative experiment in (F) to (H). Letters indicate statistical significance (ANOVA with post hoc Tukey or T-test [in (A)], *P* < 0.05).

We used RNA-seq to characterize the effect of NAM treatment, and thus overall TIR activity, on defense. We infected Col-0 and *eds1* plants with *Pst* DC3000 EV to trigger basal defense or *Pst* DC3000 *AvrRps4* to activate TNL RPS4, with or without 50mM NAM, and we identified genes inhibited by NAM treatment (*SI Appendix*, Fig. S10 and S11, Table S2; Methods). We observed that a large number of infection-regulated genes are affected by NAM (*SI Appendix*, Fig. S10A). NAM treatment alone regulated mostly genes related to “stress” or “response to chemical” (*SI Appendix*, Fig. S11). More than half of the genes regulated by NAM treatment alone were also regulated by infection (*SI Appendix*, Fig. S10B).

We defined genes regulated by infection with each strain and then subdivided these genes into NAM-sensitive (genes differentially expressed in Col-0 without NAM, but not in the presence of NAM); EDS1-dependent

(genes differentially expressed in Col-0, but not in *eds1*); and TNL RPS4-dependent genes (genes differentially expressed in Col-0 infected with *Pst* DC3000 *AvrRps4* but not with *Pst* DC3000 EV). We found that 85% and 67% of NAM-sensitive genes were also either EDS1-dependent or specifically RPS4-regulated, respectively, during *Pst* DC3000 *AvrRps4* infection (*SI Appendix*, Table S2). NAM fully inhibited 20% of genes upregulated during *Pst* DC3000 *AvrRps4* infection, but up to 41% of strictly RPS4-dependent genes (*SI Appendix*, TableS2). We conclude from these analyses that NAM predominantly affects TIR-dependent transcriptional outputs.

We hypothesized that general activation of TNLs and TIR domain proteins could also contribute to CNL-dependent immune responses. We tested the impact of NAM on CNL-dependent signaling. RPM1 and ZAR1 activate defense in response to *AvrRpm1* and *HopZ1a*, respectively, in a Ca^2+^-dependent manner and independently of EDS1 or RNLs (9, 46). 50mM NAM inhibited RPM1- or ZAR1-dependent growth restriction of *Pst* DC3000 *AvrRpm1* and *Pst* DC3000 *HopZ1a*, respectively, in Col-0 and in mutant plants lacking the cognate CNLs (Fig. 5D and E). In addition, NAM treatment delayed cell death induction by the TNL RPS4 and the CNL RPM1, but not by the CNL ZAR1 (Fig. 5F to H). Overall, inhibition of TIR enzymatic activity by 50mM NAM inhibits NLP20 signaling, basal defense against virulent bacteria, and both RNL-dependent immune responses triggered by TNLs and RNL-independent immune responses triggered by CNLs. These data collectively argue for a broad role for TIR activity in defense responses.

SADR1 is required for the ADR1-L2 DV activation mimic phenotype (Fig. 1 and 3, *SI Appendix*, Fig. S2) suggesting that SADR1 is also required for ADR1-L2-driven calcium influx (47). We investigated the requirement for TIR activity on NLR-dependent Ca^2+^ influx via inhibition with 50mM NAM. We inoculated Arabidopsis expressing the [Ca^2+^]_cyt_ reporter GCamP6 (48), with *Pst* DC3000 EV (OD_600_ = 0.2), or expressing either *AvrRps4, AvrRpm1* or *HopZ1a* in the presence or absence of 50mM NAM and quantified green fluorescence as a measure of [Ca^2+^]_cyt_ (Fig. 6) (48). We found that *AvrRps4, AvrRpm1* and *HopZ1a* all triggered Ca^2+^ influx, as previously described (9, 46, 47). NAM inhibited the elevated [Ca^2+^]_cyt_ associated with *Pst* DC3000 *AvrRps4, AvrRpm1* or *HopZ1a* infection (Fig. 6). Interestingly, *HopZ1a* induced the highest [Ca^2+^]_cyt_ levels and NAM treatment reduced this to levels similar to *AvrRpm1* treated samples, consistent with the differential impact of NAM on *AvrRpm1* and *HopZ1a*-driven cell death (Fig. 5). Overall, these results suggest that TIR enzymatic activity, as revealed by NAM inhibition, is required for increased [Ca^2+^]_cyt_ levels in various NLR-activation contexts.

**Figure 6.**
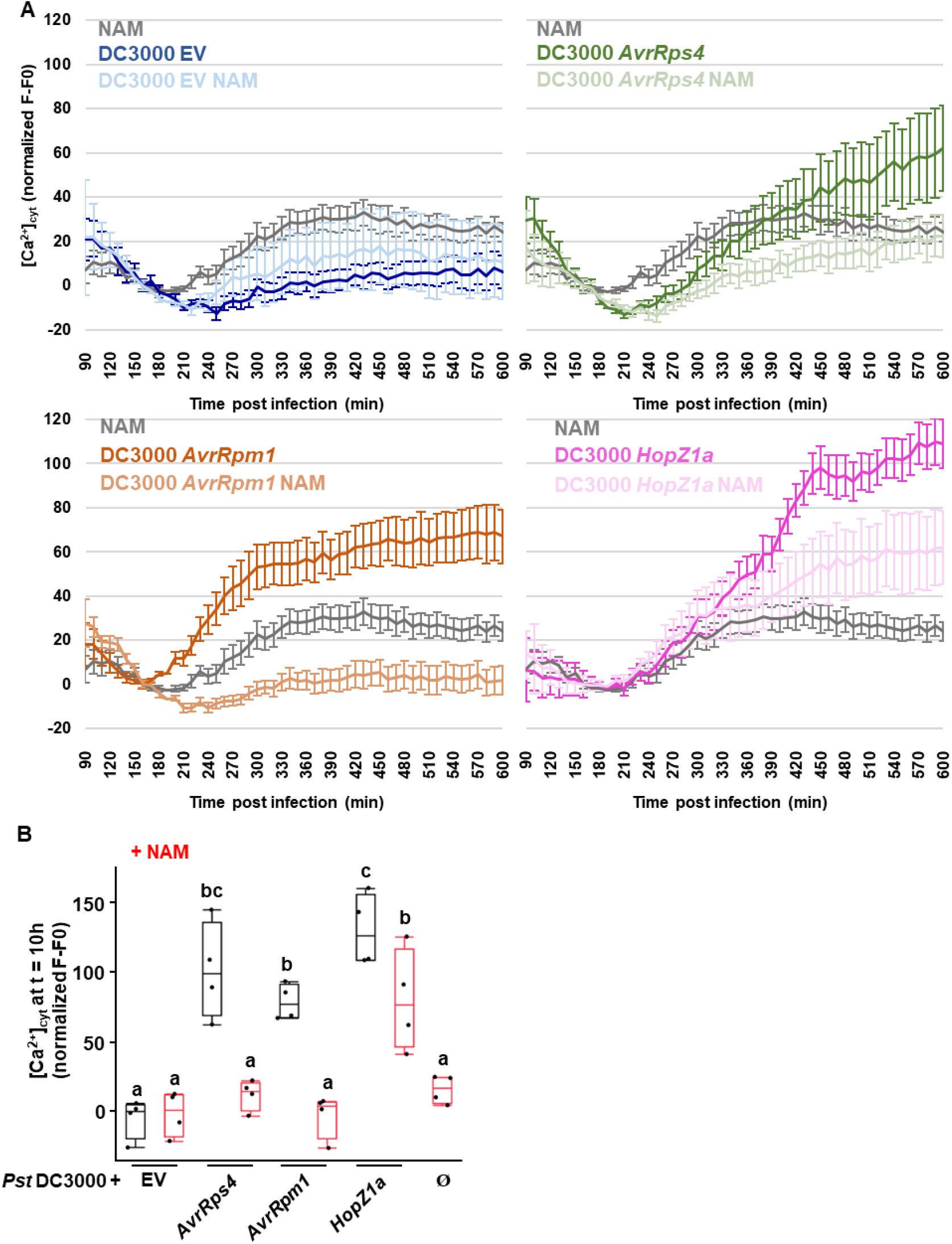
Impact of NAM 50mM on cytosolic calcium elevation during NLR mediated ETI. (**A**) Impact of NAM on [Ca^2+^]_cyt_ during ETI triggered by *Pst* DC3000 EV, *Pst* DC3000 *AvrRps4* (triggering TNL RPS4), *Pst* DC3000 *AvrRpm1* (triggering CNL RPM1) and *Pst* DC3000 *HopZ1a* (triggering CNL ZAR1), inoculated at OD_600_ = 0.2. Bars represent SEM (*N* = 8). Data from the four panels is from a single experiment and has been split into four for clarity. (**B**) Data from four experiments showing [Ca^2+^]_cyt_ at 10h post inoculation. Letters indicate statistical significance (T-test, *P* < 0.05).

## Discussion

SADR1 is a newly defined TNL that is required downstream of RNL activation for full NLP20 response, *lsd1* runaway cell death and *PR1* expression at the border of a bacterial inoculation site (Fig. 2). These three phenomena each involve a spatial component (*23*). We hypothesize that this pattern could be explained by non-autonomous signaling from pathogen-engaged to non-engaged cells, as in *lsd1* (15) or by a gradient of effector triggered-defense inhibition in neighboring cells. The cells expressing *PR1* during the response to *Pst* DC3000 *AvrRpt2* are not dead at the time of observation, contrary to the cells directly in contact with the bacteria, and therefore are likely not activating RPS2 (31). In addition, infection with *Hpa* Emwa1, which triggers the TNL RPP4, leads to strong *PR1* expression in a layer of cells bordering the cells in direct contact with the oomycete (30). NLR signaling in one dying cell would result in the leakage of immunogenic molecules, damage associated molecular patterns or reactive oxygen, activating immunity in nearby naïve cells. It is worth recalling that ADR RNLs are required for non-autonomous feed-forward cell death signaling, as *ADR1-L2* mutation suppresses runaway cell death (16). A non-autonomous immune signaling mechanism was implicated in the context of cell damage, but a requirement for RNLs was not tested (28, 29). Thus, a role in potentiating non-autonomous signaling from pathogen-engaged to non-engaged cells provides a harmonious explanation of SADR1 function.

We hypothesized that SADR1-mediated regulation of immune response could define a pathogen containment strategy. We found that *Pst* DC3000 EV could infect plants systemically in the absence of ADR1s (Fig. 3). Consistent with these results, *Pst* DC3000 can propagate systemically in *N. benthamiana* only in the absence of TNL activation (49). RNL mutants lacking ADRs exhibited systemic symptoms (growth retardation, anthocyanin accumulation and lesions) whereas Col-0 did not (Fig. 3). Therefore, RNLs and consequent Ca^2+^ signaling limit disease from localized *Pst* DC3000 EV infection events. These results highlight the importance of bacterial containment as a disease resistance mechanism. We note that *Pst* DC3000 triggers basal defense which involves weak ETI and upregulation of TIR-domain proteins and is thus at least partially TIR-dependent (24, 42, 43, 50).

We noted that SADR1 is required for ADR1-L2 DV over-accumulation, suggesting that SADR1 is required for an as yet unidentified process ultimately affecting ADR1-L2 transcription and protein accumulation (Fig. 1D, E, and F). However, it is difficult to distinguish if the lower accumulation of ADR1-L2 DV is the cause or the consequence of *sadr1-c1* suppression of ADR1-L2 DV phenotypes. Any ADR1-L2 DV suppressor mutation would likely lead to a reduction in *ADR1-L2* mRNA and protein levels since ADR1-L2 DV triggers its own expression (16). SADR1 also functions downstream or at the level of Ca^2+^ influx (47). A positive amplification loop was postulated in TIR-domain signaling (3) and RNLs are known to trigger self-amplification through an SA-based positive feedback loop (16). Our collective data demonstrate that SADR1 is a required component of this feedback loop.

We observed that SADR1, but not EDS1, was fully required for ADR1-L2 DV auto-activity (Fig. 4). This surprising finding suggests that SADR1 function is at least partially independent of EDS1 (Fig. 1, 2, 4 and *SI Appendix*, Fig. S1). SADR1 is, however, dispensable for other TNL functions (RPS4, RPP2 and *snc1*) and is not required for ADR1-L2 function when the TNL RPS4 is activated (*SI Appendix*, Fig. S4). In addition, auto-active *snc1* partially restores ADR1-L2 DV activity in the suppressed *ADR1-L2 DV sadr1-c1* background (*SI Appendix*, Fig. S4 and S6). These results collectively suggest that the mechanism underlying SADR1 function downstream of RNLs may not be specific to SADR1 and may be shared by multiple TIR domain proteins.

ADR1-L2 is functionally redundant with the Ca^2+^-permeable channel ADR1 and possesses an N-terminal motif required for ion flux in ADR1 and NRG1.1 (11, 47). Activation of a Ca^2+^ channel should be associated with pleiotropic defects, as Ca^2+^ also regulates growth and development. However, we observed that *sadr1-c1* fully suppressed the stunted growth, defense gene expression activation mimic syndrome induced by ADR1-L2 DV (Fig. 1, 3 and *SI Appendix*, Fig. S1). Therefore, SADR1 is likely to regulate ADR1-L2 DV activity at the level of Ca^2+^ influx. Consistent with this hypothesis, inhibition of TIR NADase function with NAM decreases [Ca^2+^]_cyt_ levels in the context of both coupled TNL-RNL signaling and CNL signaling (Figure 6). We cannot rule out the possibility that NAM inhibits other NADases like Poly(ADP-ribose) polymerases or sirtuins. However, there are no other plant NADases known to regulate Ca^2+^ flux and cell death. In fact, TIRs are the only known plant NADases with ADPR cyclase activity which is linked to regulation of Ca^2+^ (51, 52). TIR enzymatic function may positively influence [Ca^2+^]_cyt_ levels by activating Ca^2+^ influx mechanisms, (including RNLs themselves), by inhibiting Ca^2+^ sequestration, or both. Interestingly, cADPR can regulate [Ca^2+^]_cyt_ levels in animals by the regulation of ryanodine receptors, a class of Ca^2+^ channels involved in the Calcium-induced Calcium Release mechanism (53). Although plants do not possess ryanodine receptors, cADPR can also regulate [Ca^2+^]_cyt_ levels in plants (54–56). TIR domains are the only known proteins with ADPR cyclase activity in plants (51). Investigating the potentially varied mechanisms by which TIRs regulate [Ca^2+^]_cyt_ is key to fully understanding the plant immune system.

## Materials and Methods

A detailed description of materials and methods used in this study can be found in the *SI Appendix*.

### Plant material and growth conditions

Plants were grown in short day conditions (8h daylength) at temperatures ranging from 21°C during the day to 18°C at night. *A*. *thaliana* mutants used in this study are in the Col-0 background. The *pADR1-L2::ADR1-L2 D484V adr1-l2-4* (16), *adr1-1 adr1-l1-1 adr1-l2-4* (25), *nrg1.1 nrg1.2* (13), *eds1-12* (34), GCaMP6 (obtained from ABRC, CS69948, (48)), RNL-free *helperless* (57), pPR1:YFP^NLS^ (31), *rps2-101C* (58), *rpm1-3* (59), *snc1* (22), *chs2-1* (60), and *zar1-3* (61) mutants have been described. The *sadr1-c1* and *sadr1-c2* mutations were introduced with CRISPR-Cas9 into *pADR1L-2:ADR1-L2 D484V adr1-l2-4* and Col-0, respectively.

### Pathogen infection assays

*Pseudomonas syringae pv tomato* DC3000 syringe-infiltrations were performed as previously described (11). Plants were covered with a humidity dome for at least 30 minutes prior the start of the experiment to facilitate infiltration. Bacteria grown overnight on solid King’s B (KB) medium at room temperature, resuspended into 1mL of 10mM MgCl_2_ and diluted to the appropriate optical density 600nm (OD_600_) in 10mM MgCl_2_. When using nicotinamide (NAM, Sigma-Aldrich N0636), the dried NAM was directly added to the infiltration solution to a final concentration of 50mM right before infiltration to limit the potential toxicity of NAM. To determine pathogen sensitivity, four leaves from four plants were infiltrated with a 1mL insulin syringe, left to dry for two hours, and covered with a humidity dome for 24h. After 3 days, four samples consisting of four 0.5cm^-2^ leaf-discs from four different plants were gathered and ground in 1mL of distilled water. Samples were serially diluted in water and 5μL were spotted on KB medium supplemented with the appropriate antibiotics. For half-leaf pathogen propagation assays, plants were infiltrated with *Pst* DC3000 at OD_600_ = 0.001. Only half-leaf were infiltrated and humidity domes were kept for 48h after infiltration. After 10 days, infiltrated leaves were gathered, surface-sterilize for 1min in 70% ethanol and the un-infiltrated part of the leaves (starting from half a mm away from the mid vein), were dissected using a sterile razor blade and dried with Kimtech wipes. Samples consisting of four half-leaves from four plants were weighed and ground in 1mL of water, serially diluted and spotted on KB supplemented with rifampicin. For dip-inoculation assays, 14-days old plants grown through a mesh in 3-inch round pots were dipped in solutions of bacteria and Silwet L77 0.02% in 10mM MgCl_2_. Samples consisting of three to five plantlets were weighted and ground in 1mL of water, serially diluted and spotted on KB supplemented with rifampicin.

The impact of NAM on hypersensitive cell death was studied by infiltrating half leaves with a saturating solution of avirulent *Pst* DC3000 (OD_600_ = 0.2) and observing the samples with UV lamps at 6 (*AvrRpm1*) or 20 (*AvrRps4* and *HopZ1a*) hours post infection. Cell death was evidenced by high green autofluorescence and loss of red chlorophyll fluorescence.

*Hyaloperonospora arabidopsidis* infection assays were performed as described (11). *Hpa* isolate Cala2 was propagated on *eds1-12* mutants for three weeks prior to infection. Plants were grown in 3-inch round pots for 11 days before being sprayed with approximately 1mL of an *Hpa* spore solution at 50,000 spores per mL. Plants were covered with a humidity dome and spores were counted after 7 days. Plants were carefully placed in a 2mL Eppendorf tube containing 1mL of water and vigorously vortexed. Spores were counted with a hemacytometer. Approximately 10 plants were used for trypan blue staining as previously described (62). Plants were placed in lactophenol-trypan blue (10 mL of lactic acid, 10 mL of glycerol, 10 g of phenol, 10 mg of trypan blue, dissolved in 10 mL of distilled water and diluted 1:2 in ethanol right before use), at 60°C for at least an hour and then destained in chloral hydrate overnight or as required. Observations were performed on the Leica DMi8 (Leica Microsystems, Wetzlar, Germany).

## Acknowledgments

We thank Zac Newland-Smith for sample processing, Drs. Sarah R. Grant, Roger W. Innes and Jonathan D. G. Jones for critical reading of the manuscript and the Dangl lab for discussions of the work.

## Funding

Supported by National Science Foundation (Grant IOS-1758400 to J.L.D. and M.T.N.) and BBSRC grant BB/V01627X/1 to M.R.G. and HHMI. J.L.D. is a Howard Hughes Medical Institute (HHMI) Investigator. This article is subject to HHMI’s Open Access to Publications policy. HHMI lab heads have previously granted a nonexclusive CC BY 4.0 license to the public and a sublicensable license to HHMI in their research articles. Pursuant to those licenses, the author-accepted manuscript of this article can be made freely available under a CC BY4.0 license immediately upon publication.

## Notes

### Competing Interest Statement

The authors have declared no competing interest.

