## Supplementary material for "Broader functions of TIR domains in Arabidopsis immunity"

**This PDF file includes:**

Supplementary materials and methods.
Figures S1 to S11
Tables S1 to S3
Legends for Datasets S1 to S4
SI References

**Other supporting materials for this manuscript include the following:**

Datasets S1 to S4

### Materials and methods

CRISPR-cas9 editing:

pMR286 (kindly provided by Adi Avni, unpublished plasmid collection of Mily Ron) was modified as follows: the FastRed selection marker was amplified from BCJJ360A (1) and inserted instead of the kanamycin resistance cassette using HindIII to produce pMR286FR. Subsequently, the egg cell specific promoter (Ecp) and enhancer sequences from pHEE401E (2) were inserted into EcoRI-digested pMR286FR to drive Cas9 expression. A polycistronic gRNA-tRNA construct containing two gRNAs was synthesized (Genescript) and inserted into pMR286FR-Ecp with StuI and SpeI. Primers and guide RNA sequences used in this study are described in Table S3.

NGS-based bulk segregant analysis:

ADR1-L2 DV M3 EMS mutants exhibiting a wild type (WT) phenotype were backcrossed to ADR1-L2 DV and approximately 100 segregating F2 plants showing a suppressed phenotype were used for bulk segregant analysis. DNA was extracted with Qiagen DNeasy 96 (Qiagen, 69581). Libraries were made from the bulk genomic DNA using the KAPA HyperPrep kit (Roche, 07962363001) and KAPA Unique Dual-Indexed Adapter Kit (Roche 08861919702). Libraries were sequenced using Hiseq2000 (paired-end, 150bp) and reads were trimmed and aligned to the reference genome (Mismatch cost 2, insertion cost 3, deletion cost 3, length fraction 0.95, similarity fraction 0.95, TAIR10, CLC genomics V11). Variants were called and filtered for G to A and C to T transitions consistent with EMS-induced mutations. Variants of interest were located in genomic regions showing high frequency (>85%) in suppressed plants.

RNA extraction and gene expression analyses:

Total RNAs were extracted from frozen leaf tissue using RNeasy 96 (Qiagen, 74181). On-column DNase I was used to remove genomic DNA contamination (Qiagen, 79254). For Q-RT-PCRs, total RNAs were reverse transcribed

(Superscript III, ThermoFisher 18080044) using 18bp long oligo dT (Eurofins). cDNAs were diluted tenfold and used for amplification with Power SYBR Green master mix (ThermoFisher, 4367659) and gene specific primers (*SI Appendix,* Table S3).

RNA-sequencing libraries were constructed with KAPA mRNA HyperPrep Kit (Roche, 08098123702) and KAPA Unique Dual-Indexed Adapter Kit (Roche 08861919702). Libraries were sequenced on HiSeq2000 (ADR1-L2 DV and suppressor mutants, *SI Appendix,* Fig. S1, paired-ends 50bp) or NextSeq2000 (*SI Appendix,* Fig. S10, paired-ends, 150bp). Reads were trimmed and aligned to the reference genome to determine gene expression levels (RPKM) with RNA-seq analysis tool (mismatch cost 2, insertion cost 3, deletion cost 3, length fraction 0.8, similarity fraction 0.8, TAIR10, CLC genomics V11). Differential expression for RNA-seq was used to determine significantly misregulated genes compared to the control (Col-0 or Col-0 mock treated for Fig. S1 and S10 respectively). Differentially expressed genes were used to make heatmaps (Manhattan distance, average linkage, fold change >2) or volcano plots. Venn diagrams were made using venny 2.1 (https://bioinfogp.cnb.csic.es/tools/venny/). Gene ontology enrichment and classification was performed using PANTHER V17 (http://www.pantherdb.org/data/) and TAIR (https://www.arabidopsis.org/tools/bulk/).

Bioinformatic analyses:

Phylogenetic tree was built from Arabidopsis NB-ARC (PF00931) containing protein sequences selected using HHMER v3.3.2 (-E 1 --domE 1 --incE 0.01 --incdomE 0.03 --seqdb uniprotrefprot , http://hmmer.org/). The 326 sequences obtained were aligned with wasabi (http://was.bi/) with the pagan realignment tool using standard parameters. Aligned sequences were used to build a maximum likelihood phylogenetic tree using IQ-TREE (version 1.6.12, options: -nt AUTO -ntmax 5 - alrt 1000 -bb 1000, (3)). The tree was visualized with ITOL (https://itol.embl.de/, (4)).

Homology based structural modeling was made on swissmodel (swissmodel.expasy.org/) based on RPP1 crystal structure (7CRC, (5)). The model with the most accuracy and coverage was used visualized aligned to RPP1 structure using CLC drug design.

*lsd1 runaway cell death* measurements:

Plants were grown for four weeks before *runaway cell death* induction by spraying with 300 µM BTH diluted in water (Benzothiadiazole, Actiguard, Novartis, 53071), as previously described (6). Plants fresh and dry weight were determined after 2 weeks.

Epifluorescence observations:

Bacteria were tagged by the introduction of pMF440 (addgene plasmid n62550), driving mCherry expression, by tri-parental mating. Plants carrying the pPR1:YFP$^{NLS}$ transgene (7) were hand-infiltrated with the indicated fluorescent bacteria and left without a lid until observation. Fluorescence was observed with the Leica M205 FA (Leica Microsystems, Wetzlar, Germany) using YFP (Leica 10447410) or DSRED (Leica, 10447412) filter sets.

Western-Blots:

Leaf materials were flash frozen in liquid nitrogen and total proteins were extracted with extraction buffer (125 mM Tris-HCl, pH 8.8, 1% (w/v) SDS, 10% (v/v) glycerol, 50 mM Na2S2O5 and 1x plant protease inhibitor cocktail, P2714, Sigma-Aldrich, Saint Louis, MO, USA). Lysate was cleared by centrifugation at 20,000.*g* for 10 minutes at 4ºC. Protein concentration was determined with a Bradford assay (Biorad protein assay, 500-0006 Biorad, Hercules, CA, USA). Proteins were run on an 8% acrylamide gel and transferred to a PVDF membrane. Membranes were blocked with 3% non-fat milk in TBST and blotted with anti-HA rat antibody diluted 2000 times (Roche, 11867431001, Roche Holding AG, Basel, Switzerland) and then with an HRP-conjugated anti-rat antibody (ab97057, Abcam, Cambridge, UK).

Intracellular [$Ca^{2+}$] determination:

Bacteria were hand-infiltrated into leaves of GCaMP6 reporter plants, at $OD_{600}$ 0.2 (8). After the leaves were allowed to dry for 45 minutes, 8 leaf discs of 0.5$cm^{-2}$ from 2 plants were gathered for each condition. Leaf discs were placed on 200uL of distilled water, or 200uL NAM 50mM if previously NAM-treated, in a Nunclon 96 Flat Bottom Black plates (Thermo-Fisher) and allowed to equilibrate for an additional 45min. Fluorescence was then recorded using TECAN Infinite M200 Pro plate-reader, with excitation at 470 nm (7 nm 25 bandwidth) and emission recorded at 525nm (20nm bandwidth) with 20 µs integration time and 5 ms settle time. Absolute fluorescence values for each experiment were normalized to the untreated control value as (F-F0)-($F_{t160}$-$F0_{t160}$) (where F was the measured fluorescence at a given time point, F0 was the averaged measurement for mock-treated samples at each time points and $F_{t160}$ and $F0_{t160}$ were F and F0 values at time 160 minute, time at which fluorescence reached a minimal value).

Metabolite analysis in Arabidopsis.

Six to seven week old plants were hand-infiltrated with the indicated bacteria lacking both copies of *HopAM1*(9, 10) in the presence or absence of 50mM NAM. Samples consisting of six to eight leaves from four plants were harvested 12 hours post infection, flash frozen in liquid nitrogen then lyophilized. A Waters Xevo TQXS triple quadruple mass spectrometer coupled with Waters I-class UPLC was used for LC-MS/MS analysis of v-cADPRs and related compounds from both standards and plant extracts. For electrospray ionization source in positive ion mode the following conditiois were used; Capillary voltage: 800V, desolvation temperature: 600°C, desolvation gas: 1000L/hr, cone gas 150L/hr and nebulizer gas: 7 bar. MRM transitions for v-cADPRs are parent ions at m/z 542.00 and daughter ions at 136.00 and 348.02 with collision energy of 32 and 28eV respectively. MRM transitions for 2’ or 3’-O-beta-ribofuranosyladenosine (2’/3’-RFA) are; parent ion at m/z 400.10 and daughter ion at 136.00 with collision energy at 30 eV. UPLC mobile phases comprised water with 2mM ammonium acetate (A) and methanol (B). The gradient used for elution was as follows: 0-5mins, 100% A, 5-7mins, 80% A, 7-8mins, 100% B, then isocratic for 2mins at 100% B before equilibrating back to 100% A for 15mins with a flow rate of 0.2ml/min. Column used is a Waters Acquity UPLC CSH C18, 1.7um, .1x100mm. High resolution measurements are done on a Bruker MaXis II Q-TOF mass spectrometer. 14.4ug of umbelliferon was added to each sample as an internal standard. Ion counts were normalized to umbelliferon and to mock treated samples.

Transient expression in *Nicotiana benthamiana*.

DNA fragments encoding the TIR-domains of *At4G36140* (TIR-1, TIR-2) and *At4G36150* were PCR-amplified from Col-0 genomic DNA using Q5 High Fidelity Polymerase (New England Biolabs). Open reading frames corresponding to *At4G36140*$_{1\text{-}294}$ (TIR-1), *At4G36140*$_{423\text{-}608}$ (TIR-2), and *At4G36150*$_{1\text{-}242}$ were amplified and BP-cloned into pDONR207 using Gateway$^{TM}$ BP Clonase (Invitrogen) and sequence verified (Azenta). ORF fragments were transferred from pDONR207 using Gateway$^{TM}$ LR Clonase (Invitrogen) into the previously described binary vector pGWB602 (35S: HA_SAM-), which adds an N-terminal Sterile alpha motif (from SARM1) oligomerization domain (11). The polymerase incomplete primer extension (PIPE) method was used to introduce single amino acid substitutions at the catalytic E. All PCRs were performed using Q5 High-Fidelity polymerase (New England Biolabs, Ipswich MA), and constructs were sequence verified by Azenta. The HA_SAM oligomerization domain (1X HA tag-SARM1$_{478\text{-}578}$-GGGGS) is from the human protein, SARM1, and was previously described in (11). Amino acid sequences for all TIR constructs are provided in the Data S2.

*Agrobacterium tumefaciens* strain GV3101 delivering binary constructs of interest was syringe infiltrated at $OD_{600}$ = 0.80 into ~4-5-week *N. benthamiana* (*Nb*) *eds1* leaves. Viral suppressor of silencing, p19, was included at $OD_{600}$ 0.05, as previously described (12). GV3101 liquid cultures were grown overnight at 28°C in antibiotic selection (50 µg/mL rifampicin, 50 µg/mL gentamicin and 50 µg/mL spectinomycin, or 30 µg/mL kanamycin). Cultures were pelleted at 5k RPM for 5 min and resuspended in induction buffer [10 mM MES buffer (pH 5.60), 10 mM $MgCl_2$, and 100 µM acetosyringone] for ∼3 h prior to infiltration. *Nb* plants were grown in a Conviron environmental chamber set at 25°C, 70% humidity, and 16 h light [80 µE·m−2·s−1]. Cell death was observed at 4 days post infiltration. For LC-MS/MS analyses, tissues were collected at 40 h post *Agro*-infiltration.

Metabolite analysis was performed on a Waters TQ-XS triple quad mass spectrometer coupled with a Waters H-class UPLC. Column used is a Waters Acquity UPLC-C18, 1.7 µm, 2.1x50mm. Mobile phases include A: 2mM ammonium acetate and B: methanol. Flow rate: 0.2 mL/min. Gradient: 0-5 min, 0% B; 5-7min, 0% to 20% B; 7-8min, 20% to 100% B; then the column is washed with 100% B for 2 mins before equilibration to 100% A for 15 mins. Mass spectrometer conditions: Capillary voltage: 800V; desolvation temperature: 600°C; desolvation gas (nitrogen, 1000 L/hr); cone gas: 150 L/hr, Nebuliser gas: 7 bar. MRM parameters for the detection of cADPR-isomers, 542/136 (cone voltage 20V, collision energy: 32eV); 542/348 (cone voltage 20V, collision energy: 28eV). 14.4ug of umbelliferon was added to each sample as an internal standard. Ion counts were normalized to umbelliferon.

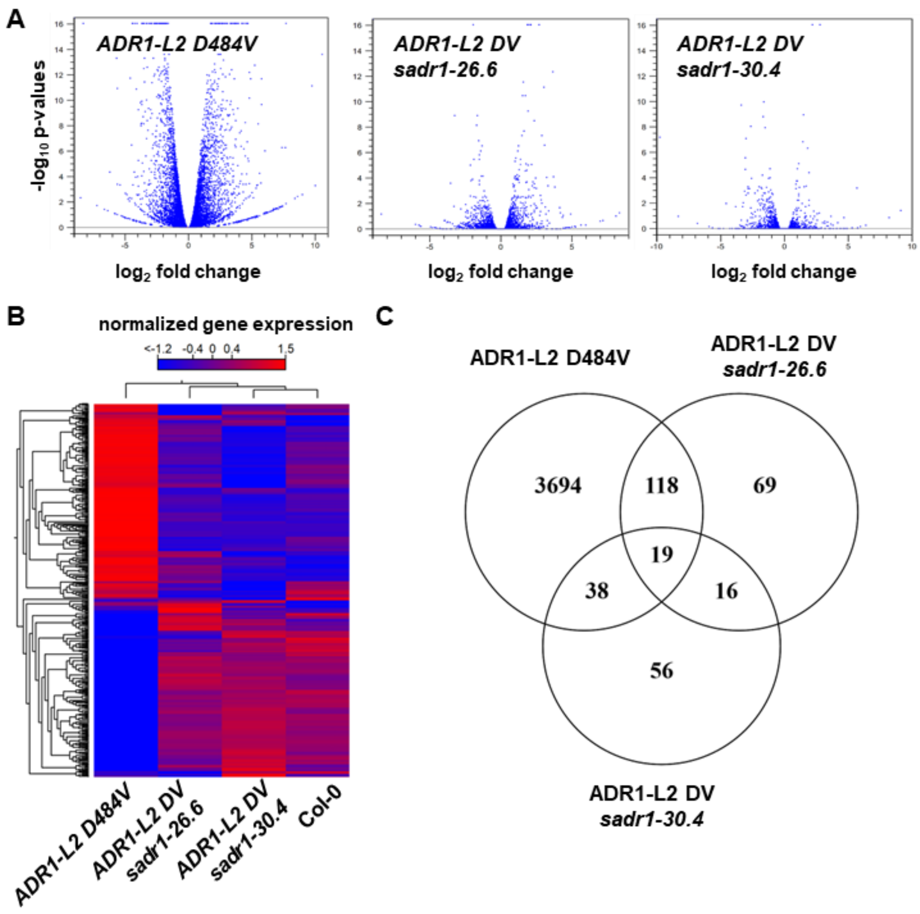


**Figure. S1. 26.6 and 30.4 mutants are full suppressors of ADR1-L2 DV induced gene expression.**

(**A**) Volcano plot showing significant gene expression changes in *ADR1-L2 DV*, *ADR1-L2 DV sadr1-26.6* and *ADR1-L2 DV sadr1-30.4* compared to Col-0 (FDR < 0.01). (**B**) Heatmap representation of gene expression levels in the different genotypes (FDR < 0.01). Interestingly, genes upregulated by ADR1-L2 DV seem to be expressed less in *ADR1-L2 DV sadr1-30.4* than in Col-0. (**C**) Venn diagram showing the overlap between misregulated genes in *ADR1-L2 DV*, *ADR1-L2 DV sadr1-26.6* and *ADR1-L2 DV sadr1-30.4,* compared to Col-0.

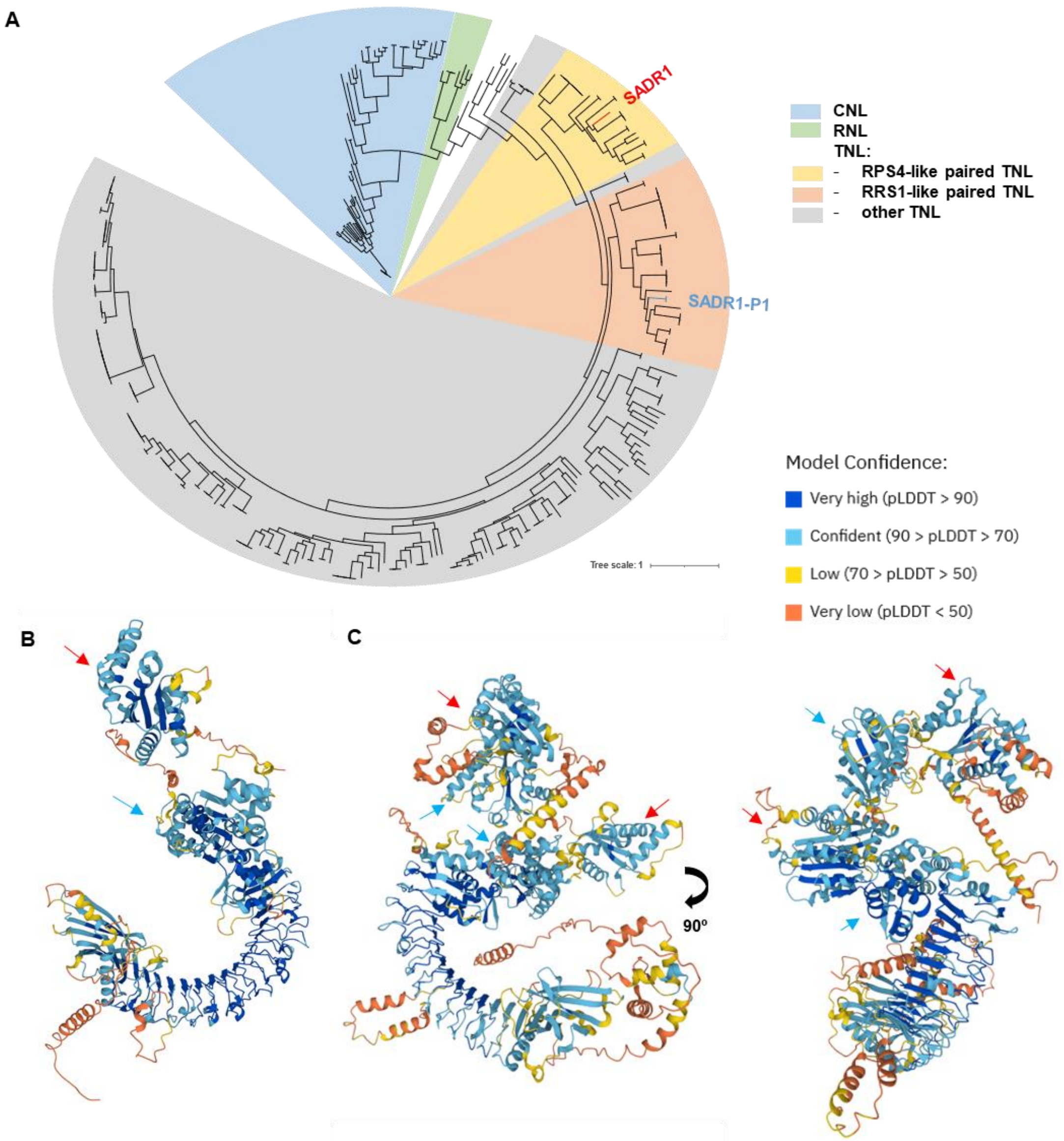


**Figure S2. SADR1 is paired with a TN-TNL.**

SADR1 (AT4G36150) and SADR1-P1 (AT4G36140) are located at the same locus, in a head-to-head configuration. (**A**) Phylogenetic tree of NB-ARC containing proteins in Arabidopsis. SADR1 is in the RPS4-like subclade of paired TNL while SADR1-P1 is present in the RRS1-like clade of paired TNL. (**B**) and (**C**) SADR1 and SADR1-P1 predicted structures (alphafold protein structure database, alphafold.ebi.ac.uk) showing TIR domains (red arrows) and NB-ARC domains (blue arrows). Note that SADR1-P1 possesses an extra N-terminal helix predicted to be transmembrane (7-26).

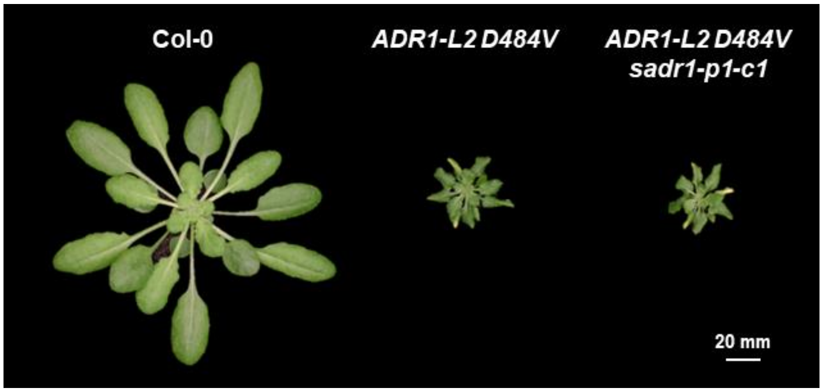


**Figure S3. SADR1-P1 is not required for ADR1-L2 D484V-driven stunted growth.**

SADR1-P1 was mutated with CRISPR. The loss of function allele *sadr1-p1-c1* (an 86bp deletion leading to a frameshift after I82 and an early stop codon after residue 202, c.247_332del) does not notably modify *ADR1-L2 D484V* auto-activity.

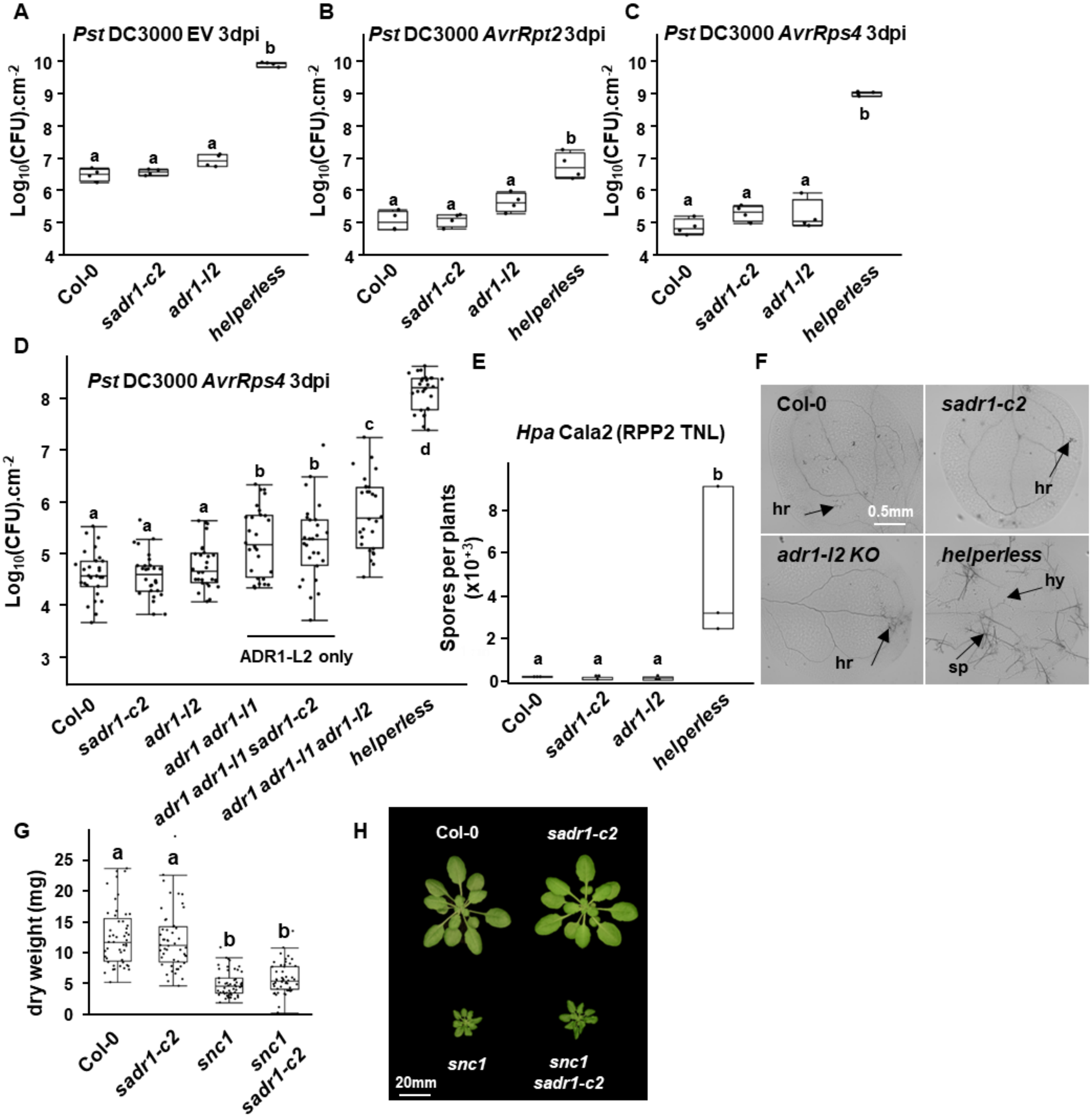


**Figure S4. SADR1 is not a helper TNL for RPS4, RPP2 and *snc1*.**

SADR1 is not required for RNL helper function during infection with *P. syringae* pv. *tomato* (*Pst*) DC3000 EV (**A**), *AvrRpt2* (**B**) or *AvrRps4* (**C**). (A), (B) and (C) show representative data from one experiment, ($N$ = 4). Experiments were performed three to four times with similar results. **(D**) SADR1 is not required for RNL helper function downstream of TNL RPS4 activation, even in the absence of the redundant ADR1 and ADR1-L1. Data presented in (D) are from seven independent experiments (*N* = 28). (**E**) SADR1 is not required for resistance against *Hyaloperonospora arabidopsidis* (*Hpa*) Cala2 which triggers the TNL RPP2 in Col-0, (*N* = 3). (**F**) Representative picture of *Hpa* Cala2 infected cotyledons stained with trypan blue (hr = hypersensitive cell death, hy = hyphae, sp = sporangiophore). (**G**) Dry weight measurements showing that SADR1 is not required for the stunted growth triggered by *snc1* TNL auto-activity, (*N* > 48). **(H**) Representative pictures of plants measured in (G). Data in (G) is from three independent experiments. Letters indicate statistical significance [ANOVA with post hoc Tukey, *P* < 0.05 or Student t-test in (E)].

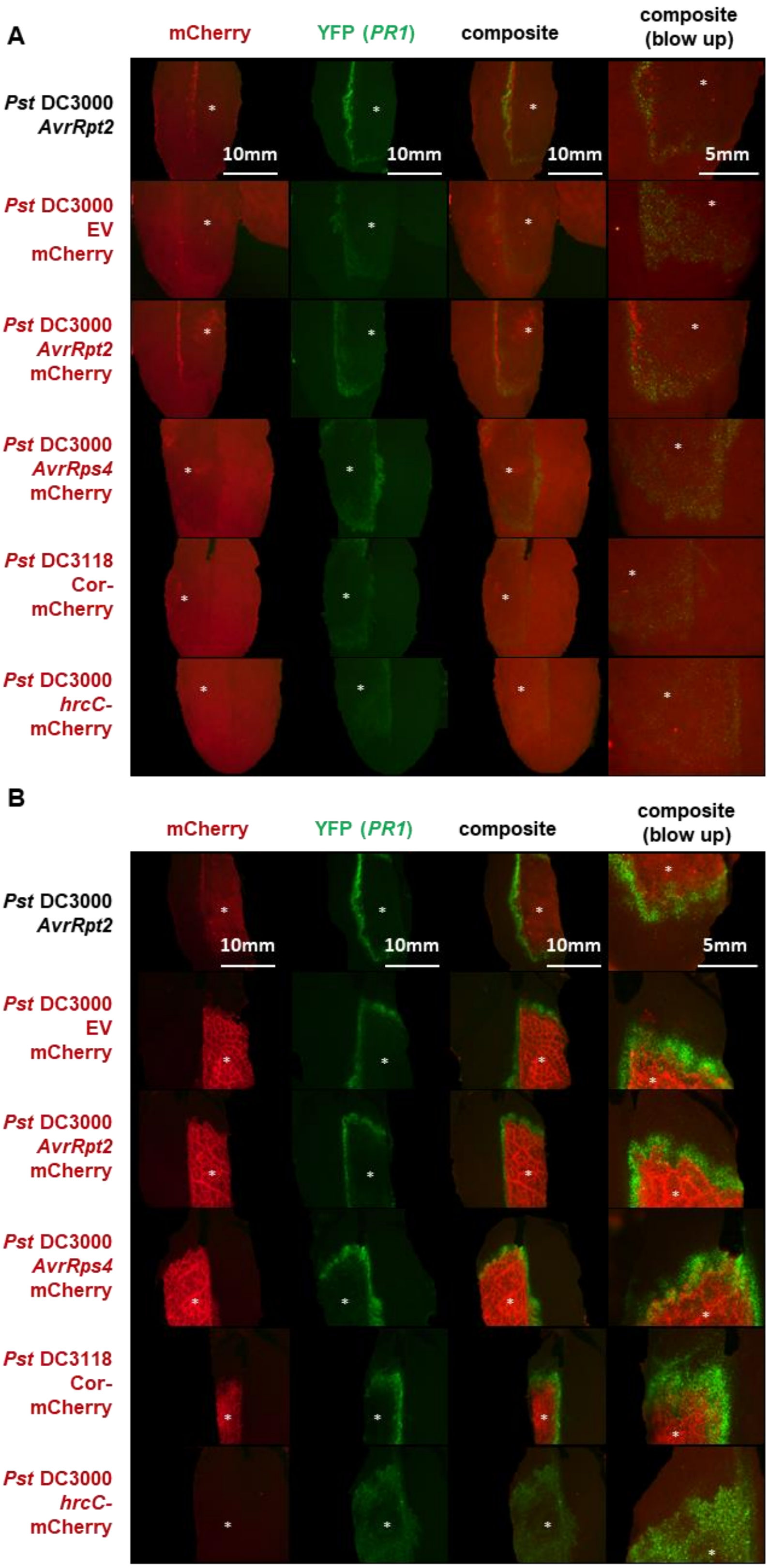


**Figure S5. Effector delivery by *Pst* DC3000 is associated with a pattern of *PR1* expression. (continued, next page)**

**Figure S5. Effector delivery by *Pst* DC3000 is associated with a pattern of *PR1* expression.**

Plants expressing nuclear-directed $YFP^{NLS}$ under the control of the *PR1* promoter were challenged with the indicated *Pst* DC3000 strains. Approximately two thirds of one side of leaves were infiltrated with the indicated bacteria at $OD_{600} = 0.2$. The infiltration site is indicated with a white asterisk. The *PR1* expression pattern is visible at 6hpi in plants infected with effector-delivering bacteria but not *hrcC*$^{-}$ mutant lacking a T3SS (**A**), suggesting that NLR activation increases gene expression on the infection border. After 24h (**B**), robust bacterial proliferation leads to a strong mCherry signal. The mCherry acquisition time is reduced in (B) to avoid signal saturation and thus chlorophyll auto-fluorescence is no longer visible. Notably, the mCherry signal and YFP signal are mostly non-overlapping, consistent with cells expressing defense genes being spatially separated from the bacterial cells.

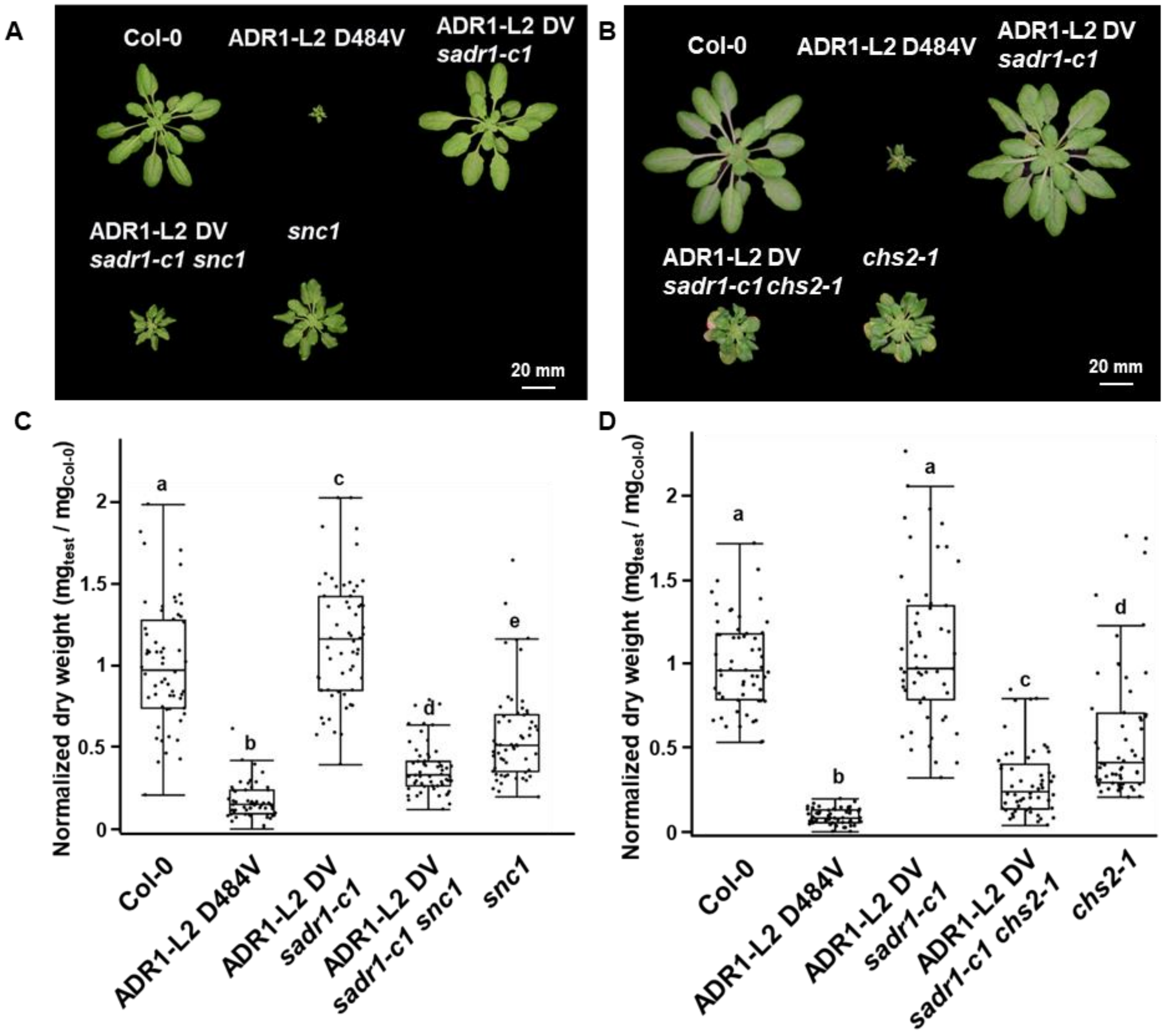


**Figure S6. The auto-active TNL alleles *snc1* and *chs2-1* partially restores ADR1-L2 DV activity in the suppressed ADR1-L2 DV *sadr1-c1*.**

(**A**) and (**B**) representative picture of six weeks old plants with the indicated phenotype. (**C**) and (**D**) Dry weight measurements, normalized to Col-0, of plants shown in (A) and (B) respectively. Data are from three independent experiments ($N$ > 48). Letters indicate statistical significance (ANOVA with post hoc Tukey, $P$ < 0.01).

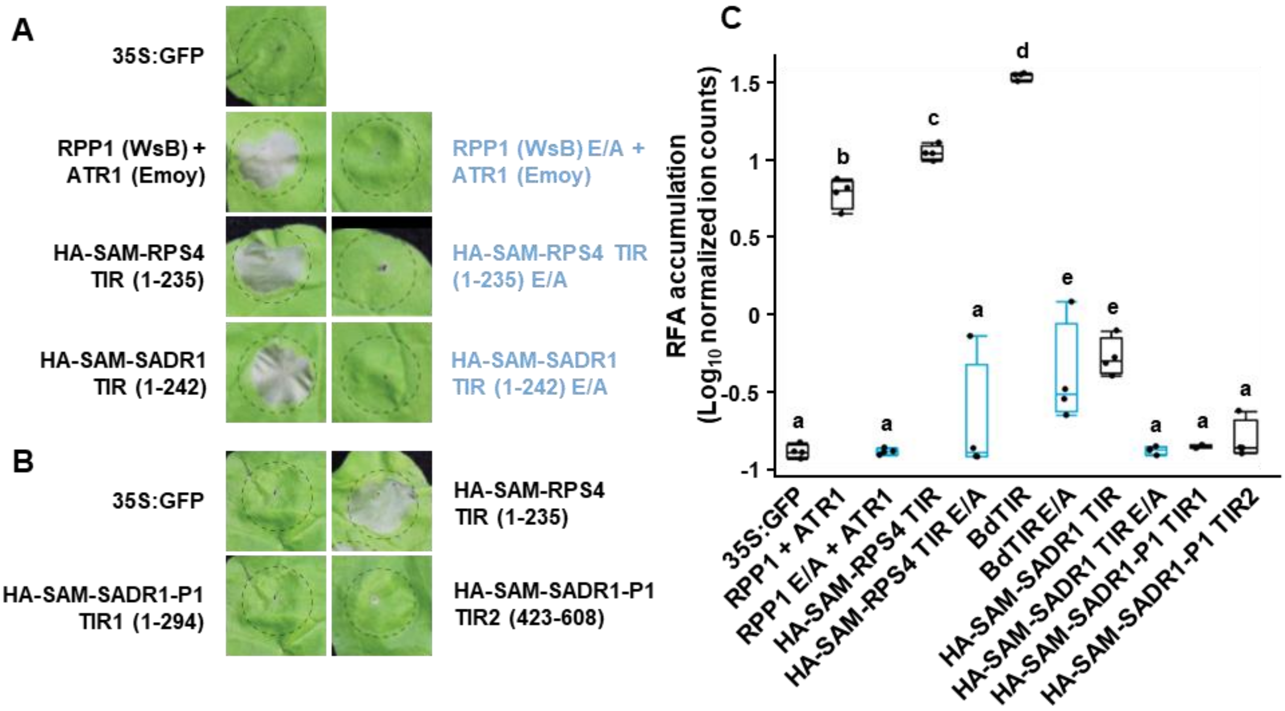


**Figure S7. Ribofuranosyl adenosine is a marker for TIR enzymatic activity.**

O-β-D-ribofuranosyl adenosine (RFA) accumulation and cell death activation by TIR domains requires the catalytic glutamic acid. (**A**) Cell death activation by various TIR domain containing proteins at four days after inoculation. Notably, the SADR1 TIR domain fused to SARM1 oligomerization domain (SAM) can trigger cell death in WT *Nic*otiana *benthamiana*. (**B**) SADR1-P1 possesses two TIR domains, but neither activates cell death. (**C**) Active TIR domains reliably trigger the accumulation of RFA in *N. benthamiana*. The accumulation is mostly dependent on the conserved catalytic E residues (in blue). Constructs were expressed in *eds1 N. benthamiana* leaves and tissues were gathered at 40hpi for RFA detection using LC-MS/MS and a 3'-O-β-D-ribofuranosyl adenosine standard. Interestingly, SADR1 TIR triggers little to none RFA accumulation. Data presented is from one representative experiment ($N = 4$, T-test, $P < 0.05$)

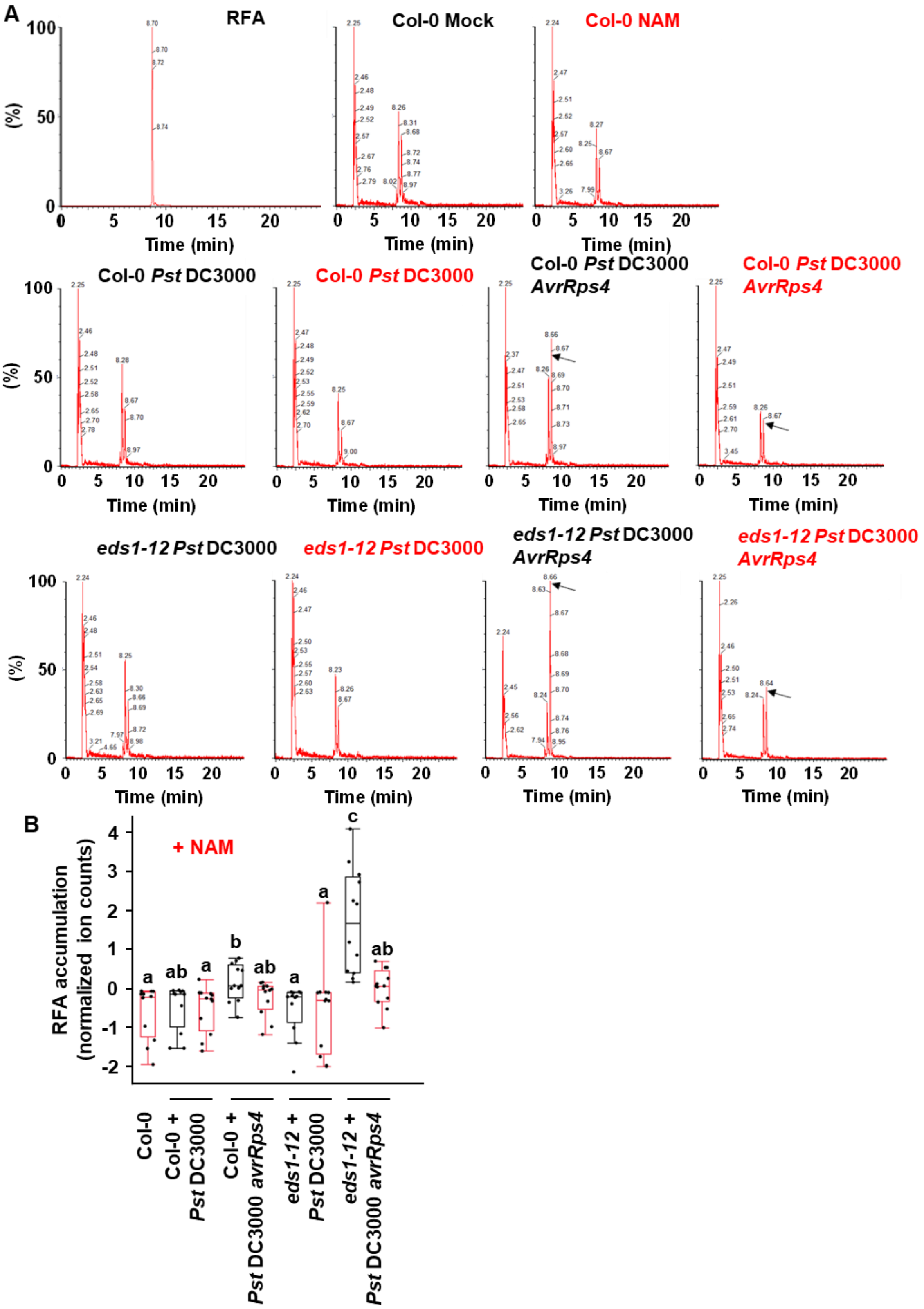


**Figure S8. NAM inhibits TIR enzymatic activity *in planta*. (continued next page)**

**Figure S8. NAM inhibits TIR enzymatic activity *in planta*.**

NAM inhibits TNL RPS4-dependent accumulation of RFA *in planta*. Plants were treated with the indicated bacteria with (red font) or without (black font) 50mM NAM and sampled at 12hpi for RFA detection using LC-MS/MS. All bacteria were *Pst* DC3000 derivative JB206 (9) which lack both copies of *HopAM1.* The experiment was performed three times with similar results. (**A**) Representative spectra from the LC-MS/MS analysis. (**B**) Samples were spiked with equal amounts of umbelliferon for relative quantification. Results are from three independent experiments ($N$ = 12, letters indicate statistical significance, $P < 0.05$, T-test).

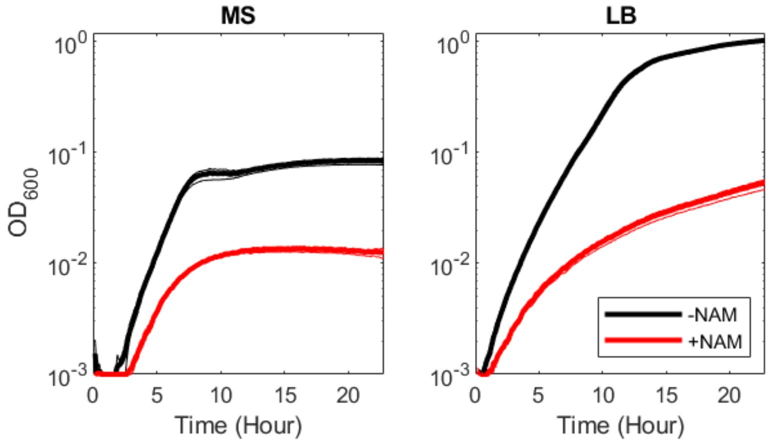


**Figure S9. 50mM NAM is bacteriostatic.**

Bacterial growth of *Pst* DC3000 as measured with $OD_{600}$ in poor (MS), or rich (LB) media supplemented or not with 50mM NAM. Importantly, this data indicates that the impact of NAM on plant defense measured in Figure 5 may be *underestimated*.

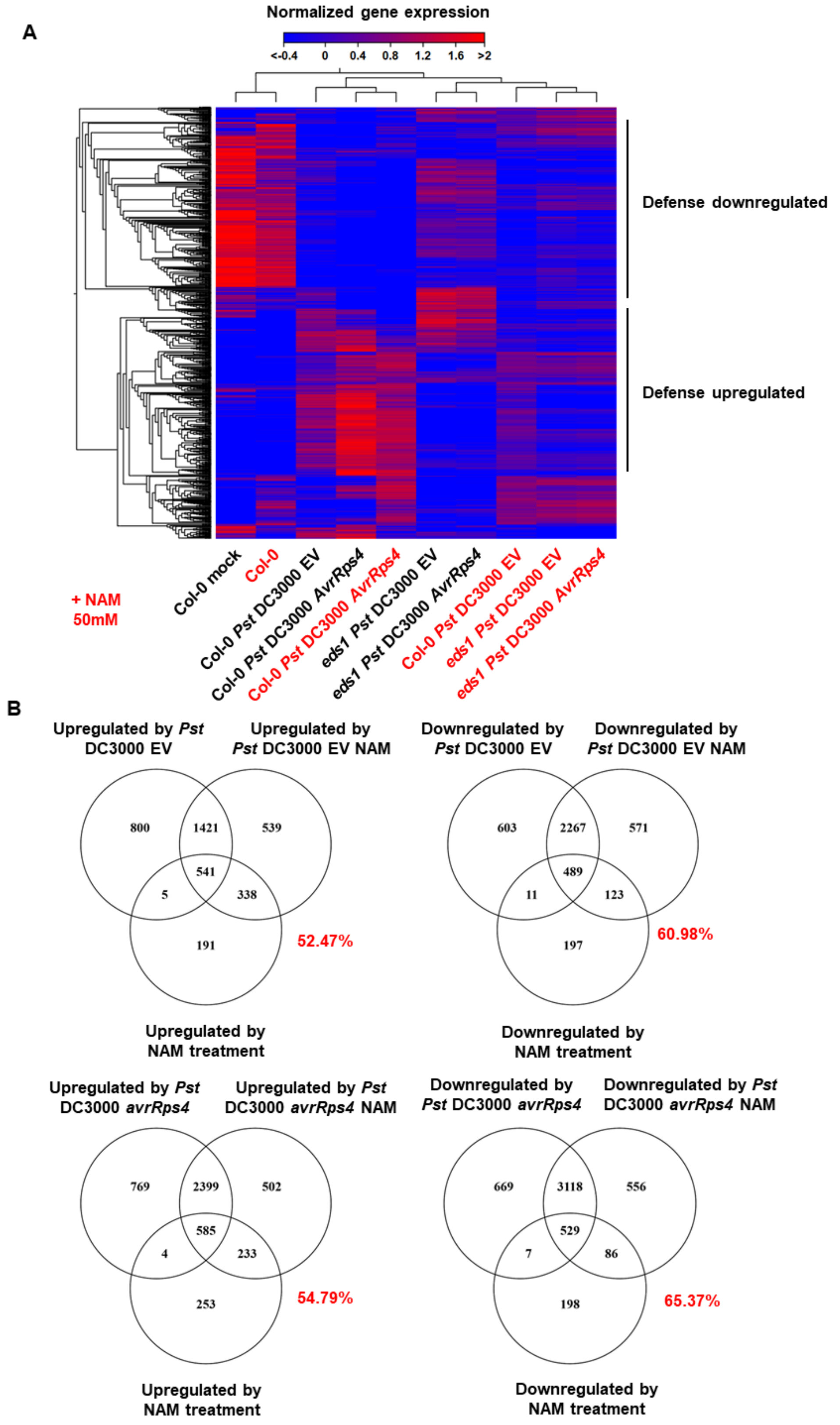


**Figure S10. NAM regulates defense. Continued.**

**Figure S10. NAM regulates defense.**

(**A**) Heatmap showing mis regulated gene expression (ANOVA, FDR < 0.01, $Log_2$ fold change > 1) upon *Pst* DC3000 EV or *Pst* DC3000 *AvrRps4* infection, with or without 50mM NAM, in Col-0 or in *eds1-12*, at 6hpi. NAM regulates the expression of a cluster of genes regardless of the genotype and treatment (NAM-regulated) and inhibits the activation of defense, in particular by *Pst* DC3000 *AvrRps4*. (**B**) Venn diagram showing the overlap between genes regulated by NAM treatment and genes regulated by pathogen infection with or without NAM treatment. Notably, NAM treatment alone mostly regulates immune regulated genes (the percentage of NAM-regulated genes also regulated by defense signaling is indicated in red). See also Table S2.

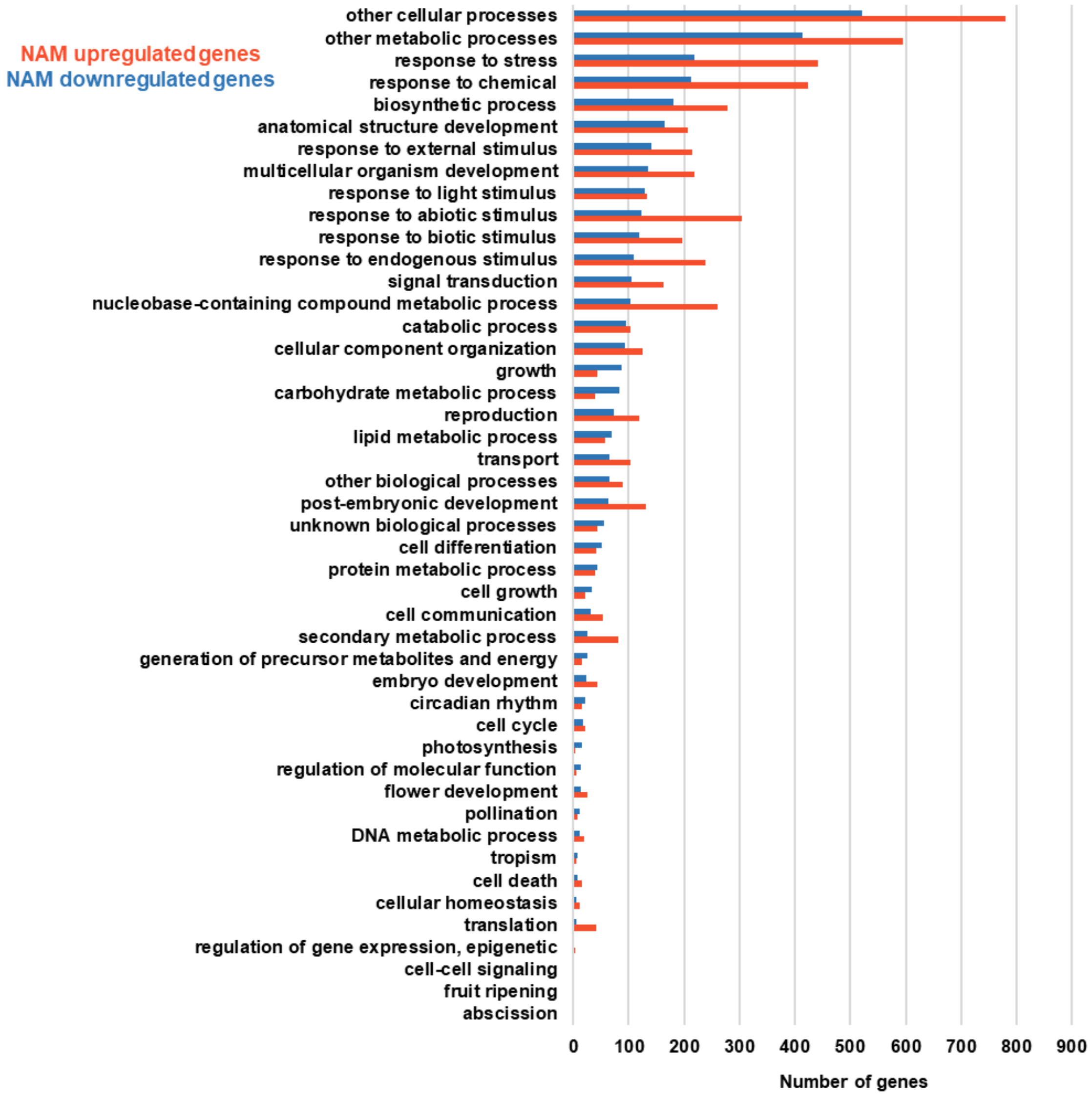


**Figure S11. Functional classification of genes upregulated by 50mM NAM treatment**

NAM treatment alone regulates the expression of 1895 genes. Most of these genes are involved in metabolic processes and stress responses.

**Table S1. GO term enriched in ADR1-L2 DV expressing plants:**
GO term enrichment analyses was performed with PANTHERV17.

| Enriched GO terms | Fold Enrichment | FDR |
|---|---|---|
| regulation of salicylic acid biosynthetic process | 10.14 | 3.69E-02 |
| response to molecule of bacterial origin | 6.26 | 2.01E-03 |
| defense response to bacterium, incompatible interaction | 5.8 | 7.67E-04 |
| regulation of jasmonic acid mediated signaling pathway | 5.32 | 9.54E-03 |
| response to chitin | 5.22 | 9.04E-14 |
| salicylic acid mediated signaling pathway | 4.94 | 4.09E-02 |
| plant-type hypersensitive response | 4.79 | 3.90E-05 |
| systemic acquired resistance | 4.74 | 8.04E-04 |
| leaf senescence | 4.35 | 6.78E-06 |
| response to hypoxia | 4.11 | 1.71E-02 |
| defense response to oomycetes | 3.91 | 1.63E-02 |
| positive regulation of defense response | 3.59 | 1.10E-03 |
| regulation of response to biotic stimulus | 3.13 | 4.86E-05 |
| positive regulation of response to external stimulus | 3.16 | 4.85E-02 |
| regulation of innate immune response | 2.99 | 2.83E-02 |
| cellular response to nutrient levels | 2.78 | 4.15E-03 |
| response to wounding | 2.55 | 8.50E-03 |
| response to jasmonic acid | 2.47 | 1.46E-02 |
| response to abscisic acid | 2.35 | 8.20E-09 |
| response to ethylene | 2.29 | 8.38E-03 |
| protein phosphorylation | 2.1 | 1.52E-10 |
| response to water deprivation | 2.09 | 3.40E-02 |
| response to osmotic stress | 1.92 | 5.39E-03 |
| hormone-mediated signaling pathway | 1.72 | 7.45E-03 |

**Table S2. NAM regulated genes show enrichment for EDS1 and RPS4-dependent genes.**

NAM and EDS1-dependency among mis-regulated genes in response to *Pst* DC3000 EV, *Pst* DC3000 *AvrRps4* and among genes upregulated by RPS4 in Col-0. RPS4-regulated genes are the genes induced in Col-0 by *Pst* DC3000 *AvrRps4* inoculation, but not by *Pst* DC3000 EV inoculation. NAM sensitive and EDS1-dependent genes are genes induced in Col-0 by *Pst* DC3000 EV or *Pst* DC3000 *AvrRps4* that are not induced upon NAM treatment or in the *eds1* mutant, respectively. NAM-sensitive RPS4-regulated genes were obtained by comparing RPS4-regulated genes to *Pst* DC3000 *AvrRps4*-regulated genes in the presence of NAM. Notably, NAM sensitive genes are enriched for EDS1 and RPS4-dependent genes (Asterisks indicate statistical significance, Hypergeometric test with Bonferroni correction, $P < 0.01$) and approximately 40% of RPS4-regulated genes are fully inhibited by NAM.

| | **Col-0 *Pst* EV-upregulated** | | **Col-0 *Pst AvrRps4*-upregulated** | | **RPS4-upregulated** | |
|---|---|---|---|---|---|---|
| | Number of genes | % of all | Number of genes | % of all | Number of genes | % of all |
| All | 2767 | 100% | 3757 | 100% | 1270 | 100% |
| NAM-sensitive | 805 | **29.09%** | 773 | **20.57%** | 520 | **40.94%** |
| EDS1-dependent | 1416 | 51.17% | 2522 | 67.13% | 1195 | 94.09% |
| NAM and EDS1-dependent | 516 | 18.65% | 657 | 17.49% | 502 | 39.53% |
| **EDS1**-dependent genes among NAM-sensitive genes | **64.10% (*)** | | **84.99% (*)** | | **96.54% (*)** | |
| **RPS4**-regulated genes among NAM-sensitive genes | NA | | **67.27% (*)** | | NA | |
| | **Col-0 *Pst* EV-downregulated** | | **Col-0 *Pst AvrRps4*-downregulated** | | **RPS4-downregulated** | |
| | Number of genes | % of all | Number of genes | % of all | Number of genes | % of all |
| All | 3370 | 100% | 4323 | 100% | 1188 | 100% |
| NAM-sensitive | 614 | **18.22%** | 676 | **15.64%** | 446 | **37.54%** |
| EDS1-dependent | 1190 | 35.31% | 2209 | 51.10% | 1102 | 92.76% |
| NAM and EDS1-dependent | 373 | 11.07% | 543 | 12.56% | 415 | 34.93% |
| **EDS1**-dependent genes among NAM-sensitive genes | **60.75% (*)** | | **80.33% (*)** | | **93.05% (ns)** | |
| **RPS4**-regulated genes among NAM-sensitive genes | NA | | **65.98% (*)** | | NA | |

### Table S3. Sequences of primers and sgRNA used in this study.

| Oligo name | Sequence (5'-3') | Use |
|---|---|---|
| p41-Fastred-f | tatgacaagcttGAGTTCTAGAATGTCGCGGAA | Cloning FastRed selection marker from bcjj360A |
| p42-Fastred-r | ttcgcaaagcttACTAAATGGAGCAACCTACTGTTTTT | |
| p332-pECpromoF | GACTAACAGAATTCtagtgaataaaagcatttgcgttt | Cloning egg cell promoter from pHEE401E |
| p333-pECpromoR | ttctcaacagattgataaggtcgaGAATTCGACTAACA | |
| gRNA-SADR1-1 | CTGCGGCGGAGATAATTGAA | CRISPR-Cas9 guides used to KO *SADR1* |
| gRNA-SADR1-2 | ATTTCCGAGGGAAACAACTG | |
| p206-Act2F | CCGCTCTTTCTTTCCAAGC | Q-RT-PCR, ACTINE 2 reference gene |
| p207-Act2R | CCGGTACCATTGTCACACAC | |
| p428-PR1-Q-F | CATACACTCTGGTGGGCCTTA | Q-RT-PCR, *PR1* |
| p429-PR1-Q-R | CGCTAACCCACATGTTCACG | |
| p337-Sadr1crisprR | tcctcgactcctggattctc | *sadr1-c2* genotyping, mutant is 394bp, WT is 444bp |
| p377-sadr1sgRNA1-F | CCATTAGCACCATTGGTTACA | |
| p11-ADR1-L2KOF | CAACGTCACCTACTTGTTTACTTCC | *adr1-l2-4* KO genotyping, amplifies mutant band at 200 bp |
| p12-ADR1-L2KOR | GTCAATTTGTTTACACCACAATATAT | |
| p13-ADR1-L2WTR | GATCATGGTGGCGAGATTTT | *adr1-l2-4* WT genotyping, amplifies WT band at 684bp |
| p30-ADR1-L2KOF2 | AGTCTAAACCATCCGCCGAC | |
| p323-lsd1Fw | CTCCAATCAGGTTGCCCATGCT | *lsd1-2* genotyping, use with p325 to amplify mutant band at ~ 700bp, with p324 to amplify WT band at ~ 470bp |
| p325-Salk_LB-a1 | ATGGTTCACGTAGTGGGCCATC | *lsd1-2* mutant genotyping, amplifies mutant band at ~ 700bp |
| p324-Lsd1Rev | CACCAACTTTCCGCTTTCATCA | *lsd1-2* mutant genotyping, amplifies WT band at ~ 470bp |
| p21-DVinsertion2F | AGCGAGAGTTGCAGGAGAGA | Genotyping of *ADR1-L2 D484V* insertion in chromosome 4, use with p22 to amplify WT band at 250bp, use p92 to amplify mutant band at 170bp |
| p22-DVinsertion2R | TTGTGAATCCGGTTAATCAAAA | Genotyping of *ADR1-L2 D484V* insertion in chromosome 4, gives 250bp if WT |
| p92-DVinsert2R-2 | ccgcaatgtgttattaagttgtc | Genotyping of *ADR1-L2 D484V* insertion in chromosome 4, gives 170bp if mutant |
| LJ_79 | CAAAGGACGATGATGTTCGAG | genotyping *ADR1* locus, gives roughly 850bp if WT, courtesy of Lance Jubic |
| LJ_80 | CGGATTGTTCACTATAGTAAGG | |
| LJ_81 | TTTCATAACCAATCTCGATACAC | genotyping *adr1-1*, gives roughly 600bp if mutant, courtesy of Lance Jubic |
| LJ_82 | CGGATTGTTCACTATAGTAAGG | |
| LJ_83 | ATGGCCATCACCGATTTTTTC | genotyping *ADR1-L1* locus, gives roughly 900bp if WT, courtesy of Lance Jubic |
| LJ_84 | GTCAGGAACAGGATTTCCAG | |
| LJ_85 | TTTCATAACCAATCTCGATACAC | genotyping *adr1-l1-1*, gives roughly 1kb if mutant, courtesy of Lance Jubic |
| LJ_86 | GTCAGGAACAGGATTTCCAG | |
| p344-pBGYN-F | ACATGGTCCTGCTGGAGTTC | genotyping *pPR1:YFPNLS*, primers are specific to pBGYN binary vector, gives 474bp if mutant |
| p27-M13Frev | gtaaaacgacggccagt | |
| p345-JS959 | AACTAGCATACAGAGGGGCA | Genotyping *eds-12*, from (13), gives 550bp if mutant |

### Table S3. Sequences of primers and sgRNA used in this study. Continued.

| | | |
|---|---|---|
| p346-JS960 | GCTGAGAGAAATCGAACCGG | Genotyping *eds1*-12, from (13), use with p345 to amplify mutant band at 550bp or with p347 to amplify WT band at 700bp |
| P347-JG08 | aaagaagacaacattGATCTATATCTATTCTCTTTTCTT | Genotyping *eds-12*, from (13), gives 700bp if WT |
| p411-egfp-f | CATGGTCCTGCTGGAGTTCGTG | genotyping of *GCaMP6*, primers amplify a part of *eGFP*, gives 408bp if mutant |
| p412-exfp-r | GTCTTGTAGTTGCCGTCGTC | |
| p14-snc1-dcapsf | ttaaaaagggttgcgattgtt | DCAPS genotyping of *snc1*: cut PCR product with XmnI, WT band is 105bp, mutant band is 127bp |
| p15-snc1dcapsr | ggtagatccccgtaatGaAcaattt | |
| p156-26_6DCAPSF | TGTGCTGTTGTCACATTCCCT | DCAPS genotyping of *sadr1-26.6*: cut PCR product with Hpy1888III, uncut is 162bp, WT band is 110bp, mutant band is 134bp |
| p98-26_6DCAPSR | AAACACATGATCTGAATCAATCTTT | |
| p93-30_4DCAPSR | CGCAACTGTTTTCAGTTTTTCGCGG | DCAPS genotyping of *sadr1-30.4*: cut PCR product with HaeIII, uncut is 288bp, WT band is 168bp, mutant band is 193bp |
| p94-30_4DCAPSF | TTGACGGGACAGCACTTAAA | |
| p350-DCAPs-nrg1.1-2-F | CGTTGCAATGGATTCCCAATC | DCAPS genotyping of *nrg1.1 nrg1.2* double mutant from (1): cut PCR product with AcuI, WT band is 120bp + 330bp, mutant band is uncut at 450bp |
| p351-DCAPs-nrg1.1-2-R | AAACCGTCTTCGTGCTCATT | |
| p409-sadr1c1-f | GTCTTCCTCAATTTCCGAGGGAAACAg | DCAPS genotyping of *sadr1-c1*: cut PCR product with PvuII, WT band is 303bp, mutant band is uncut at 327bp |
| p410-sadr1c1-r | aacgtgtaaccaaaagcgcc | |

**Dataset S1. Gene expression regulation by ADR1-L2 D484V in Col-0 and suppressed mutants (separate file).**

Gene expression values (RPKM) and differential expression in ADR1-L2 D484V, ADR1-L2 DV *sadr1-26.6* and ADR1-L2 DV *sadr1-30.4* compared to Col-0, used for Figure S1.

**Dataset S2. SADR1 and SADR1-P1 TIR sequences (separate file)**

DNA and amino acid sequences corresponding to the construct used in Figure S7.

**Dataset S3. Gene expression regulation by NAM (separate file).**

Gene expression values (RPKM) and differential expression in Col-0 or *eds1-12*, infected or not with *Pst* DC3000 EV or *AvrRps4*, in the presence or absence of NAM, used for Figure S10 and S11.

**Dataset S4. Lists of genes regulated by NAM treatment or by infection with or without NAM (separate file).**

List of genes used for analyses in Figure S10B and Table S2.
